# Cryo-EM structure of the bicarbonate receptor GPR30

**DOI:** 10.1101/2024.05.19.594840

**Authors:** Shota Kaneda, Airi Jo-Watanabe, Hiroaki Akasaka, Hidetaka S. Oshima, Takehiko Yokomizo, Wataru Shihoya, Osamu Nureki

**Affiliations:** Department of Biological Sciences, Graduate School of Science, The University of Tokyo, Bunkyo, Tokyo 113-0033, Japan; Department of Ion Signaling and Response, The Sakaguchi Laboratory, Keio University School of Medicine, Shinanomachi, Tokyo 160-0016, Japan; Department of Signal Exploration, The Sakaguchi Laboratory, Keio University School of Medicine, Shinanomachi, Tokyo 160-0016, Japan; Department of Biochemistry, Juntendo University Graduate School of Medicine, Tokyo 113-8421, Japan

## Abstract

G-protein-coupled receptor 30 (GPR30) is a bicarbonate receptor that plays a vital role in cellular responses to extracellular pH and ion homeostasis. Despite its significance, the mechanisms by which GPR30 interacts with bicarbonate ions remain elusive. There is no consensus on a drug that targets GPR30, and difficulties in pharmacological analyses have limited biological and drug discovery research on GPR30. Here, we present the cryo-electron microscopy structure of human GPR30 in the presence of bicarbonate ions, at 3.15 Å resolution. Our structure reveals unique extracellular pockets and critical residues for bicarbonate binding and activation. Functional assays demonstrate that mutations in these residues impair bicarbonate-induced GPR30 activation, underscoring their importance in receptor function. This study also provides insights into G-protein coupling, highlighting the structural divergence between GPR30 and other GPCRs. Our findings not only advance the understanding of the role of GPR30 in pH homeostasis but also pave the way for the development of high-affinity drugs targeting GPR30 for therapeutic interventions in diseases associated with acid-base imbalance.

## Introduction

The extracellular environment regulates cellular responses, as exemplified by pH and ion homeostasis. The acid-base balance, largely based on the bicarbonate buffer system *in vivo*, serves as a vital mechanism that maintains the optimal pH for cellular function^1^. Changes in the extracellular pH and ion homeostasis are monitored by membrane channels and receptors, such as G protein-coupled receptors (GPCRs). However, it remains unknown whether and how the acid–base balance-related ions, protons and bicarbonate ions, bind to the receptors and cause the conformational changes that lead to intracellular signal transduction. These proton-sensing GPCRs are thought to be involved in pH homeostasis, particularly in the acidic tumor microenvironment, at inflamed sites, and during ischemia-reperfusion injury^2^. In recent years, the determination of several proton receptor structures has begun to elucidate the molecular mechanisms underlying proton sensing^3–7^.

We recently reported that a physiological concentration of bicarbonate ions, the counterpart of protons in the bicarbonate buffer system, activates G protein-coupled receptor 30 (GPR30), which leads to G_q_-coupled calcium responses^8^. Our study also demonstrated that GPR30 in brain mural cells regulates blood flow and ischemia–reperfusion injury. GPR30 was identified as a G protein-coupled estrogen receptor that mediates the rapid non-genomic action of estradiol (E2)^9^. However, despite numerous reports on the pleiotropic functions of GPR30 *in vivo*^10,11^, controversy remains regarding the responses of GPR30 to E2 *in vitro*^12^, *ex vivo*^13^, and *in vivo*^14^. The broad expression of GPR30^15,16^, including blood vessels^17,18^, stomach, and lung, has also raised the possibility of its non-estrogenic functions. We recently demonstrated that three amino acids, E115^2.61^, Q138^3.33^, and H307^7.36^, are essential for the bicarbonate-induced activation of GPR30, according to the public homology model (https://gpcrdb.org/)^19^. However, we did not examine whether bicarbonate ions interact with GPR30 and cause conformational changes. Moreover, to date, there is no consensus on a high-affinity drug that targets GPR30, and difficulties in pharmacological analyses have limited biological and drug discovery research on GPR30. To elucidate the bicarbonate–GPR30 interaction and to establish the first structural identification of an acid-base balance-related GPCR, we report the cryo-electron microscopy (cryo-EM) structure of human GPR30 in the presence of bicarbonate ions.

## Results

### Overall structure

For the structural study, we used the full-length human GPR30 sequence. To efficiently purify the stable GPCR-G-protein complex, the receptor and mini-G_sqi_ were incorporated in a ‘Fusion-G system’ by combining two complex stabilization techniques^20,21^ (Figure 1—figure supplement 1A, B). The modified receptor and G-protein were co-expressed in Sf9 cells and purified by Flag affinity chromatography in the presence of 200 mM NaHCO_3_. After an incubation with nanobody 35 (Nb35) and single chain scFv16, which binds to mini-G_sqi_, the complex was purified by size exclusion chromatography (Figure 1—figure supplement 1C, D). The structure of the purified complex was determined by single-particle cryo-EM analysis, with an overall resolution of 3.15□Å (Table 1, Figure 1—figure supplement 2, and “Methods”). As the extracellular portion of the receptor was poorly resolved, we performed receptor focused refinement, yielding a density map with a nominal resolution of 3.18 Å, which was combined with the refined map focused on G-protein. The resulting composite map allowed us to precisely build the atomic model of the components, including the receptor (residues 51 to 196, 207 to 288, and 299 to 340, G-proteins, and scFv16) (Figure 1A, B). Nb35 was not visible in the cryo-EM map, probably because of the effect of the mini-G_s_ modification, although the interactive residues with Nb35 were not changed.

**Figure 1.**
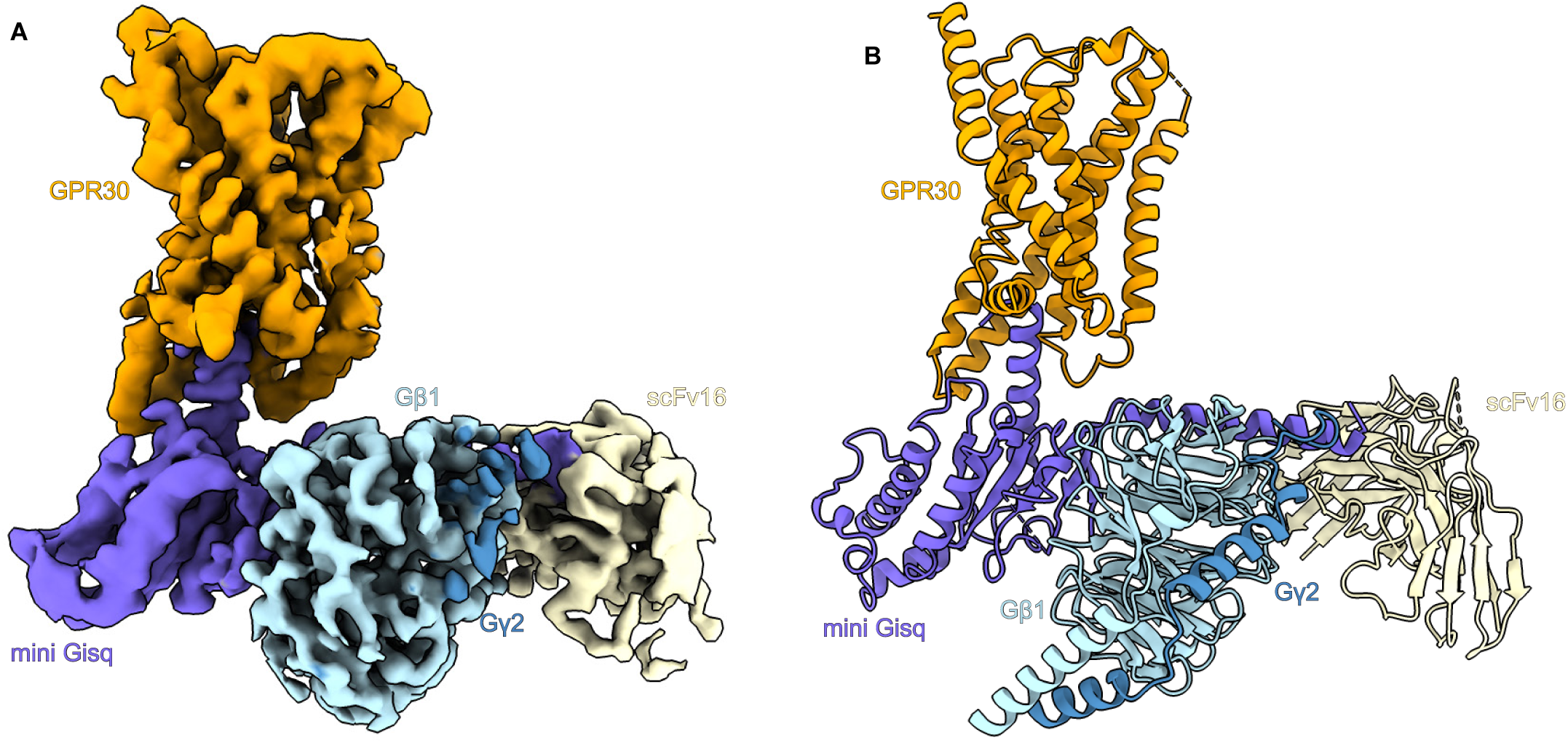
Overall structure of the GPR30-miniG_sqi_β_1_γ_2_-scFv16 complex. **A** Unsharpened cryo-EM density map of the GPR30-miniG_sqi_β_1_γ_2_-scFv16-Nb35 complex, with the components individually colored. **B** The refined structure of the complex is shown as a ribbon representation.

**Table. 1.**
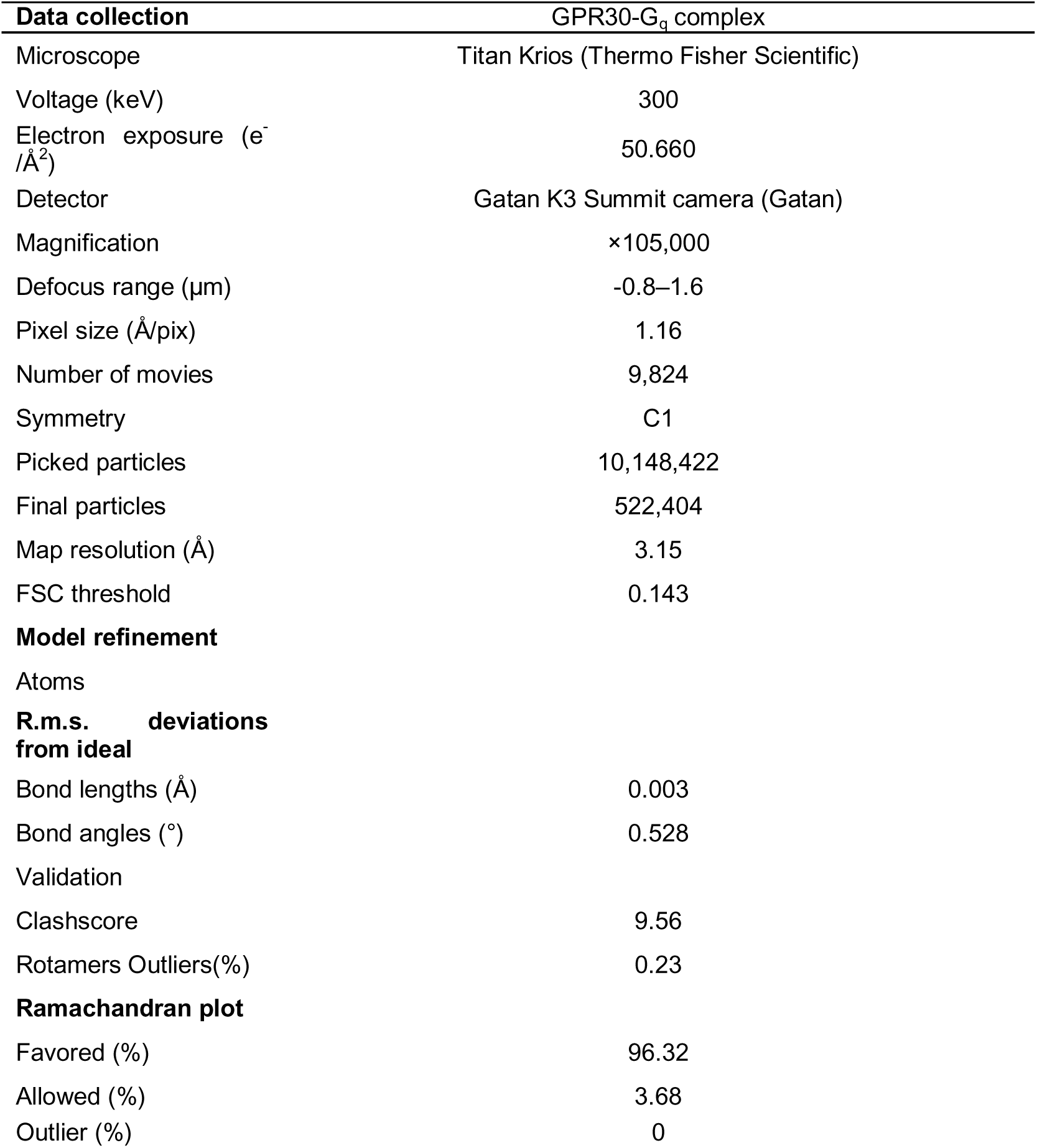
Cryo-EM data collection, refinement and validation statistics.

| **Data collection** | **GPR30-G<sub>q</sub> complex** |
| --- | --- |
| Microscope | Titan Krios (Thermo Fisher Scientific) |
| Voltage (keV) | 300 |
| Electron exposure (e<sup>-</sup>/Å<sup>2</sup>) | 50.660 |
| Detector | Gatan K3 Summit camera (Gatan) |
| Magnification | ×105,000 |
| Defocus range (μm) | -0.8–1.6 |
| Pixel size (Å/pix) | 1.16 |
| Number of movies | 9,824 |
| Symmetry | C1 |
| Picked particles | 10,148,422 |
| Final particles | 522,404 |
| Map resolution (Å) | 3.15 |
| FSC threshold | 0.143 |
| **Model refinement** |  |
| Atoms |  |
| **R.m.s. deviations from ideal** |  |
| Bond lengths (Å) | 0.003 |
| Bond angles (°) | 0.528 |
| Validation |  |
| Clashscore | 9.56 |
| Rotamers Outliers(%) | 0.23 |
| **Ramachandran plot** |  |
| Favored (%) | 96.32 |
| Allowed (%) | 3.68 |
| Outlier (%) | 0 |

The receptor consists of the canonical 7 transmembrane helices (TM) connected by three intracellular loops (ICL1–3) and three extracellular loops (ECL1–3), and the amphipathic helix 8 at the C-terminus (H8) (Figure 2A). Most of the TMs are kinked, as in typical GPCRs^22^. It should be noted that TM1 is also kinked at P71^1.44^ (superscripts indicate Ballesteros-Weinstein numbers^23^), whereas TM1 adopts a straight α-helix in most class A GPCRs. ECL1 contains a short α helix (Figure 2B). The short ICL3 was completely visible in the cryo-EM map (Figure 2C and Figure 1—figure supplement 2), while residues 197 to 206 in ECL2 and 289 to 298 in ECL3 were disordered (Figure 2B). ECL2 is attached to TM3 by the disulfide bond between C130^3.25^ and C207^ECL^^2^ (Figure 2B, Figure 2—figure supplement 1A-F), which is highly conserved in class A GPCRs^22,24^. The cryo-EM structure did not superimpose well on the AlphaFold-predicted structure (Q99527 in the AlphaFold database)^25^, with a root mean square deviation (R.M.S.D.) of Cα atoms of 3.10 □ (Figure 2—figure supplement 2A). The conserved D^3.49^R^3.50^Y^3.51^ motif in the predicted structure represents an inactive state (Figure 2—figure supplement 1B)^22^. Moreover, ECLs 2 and 3 are rich in cysteine residues, which form incorrect disulfide bonds in the predicted structure (Figure 2—figure supplement 2C). These comparisons support the usefulness of experimental structural determination.

**Figure 2.**
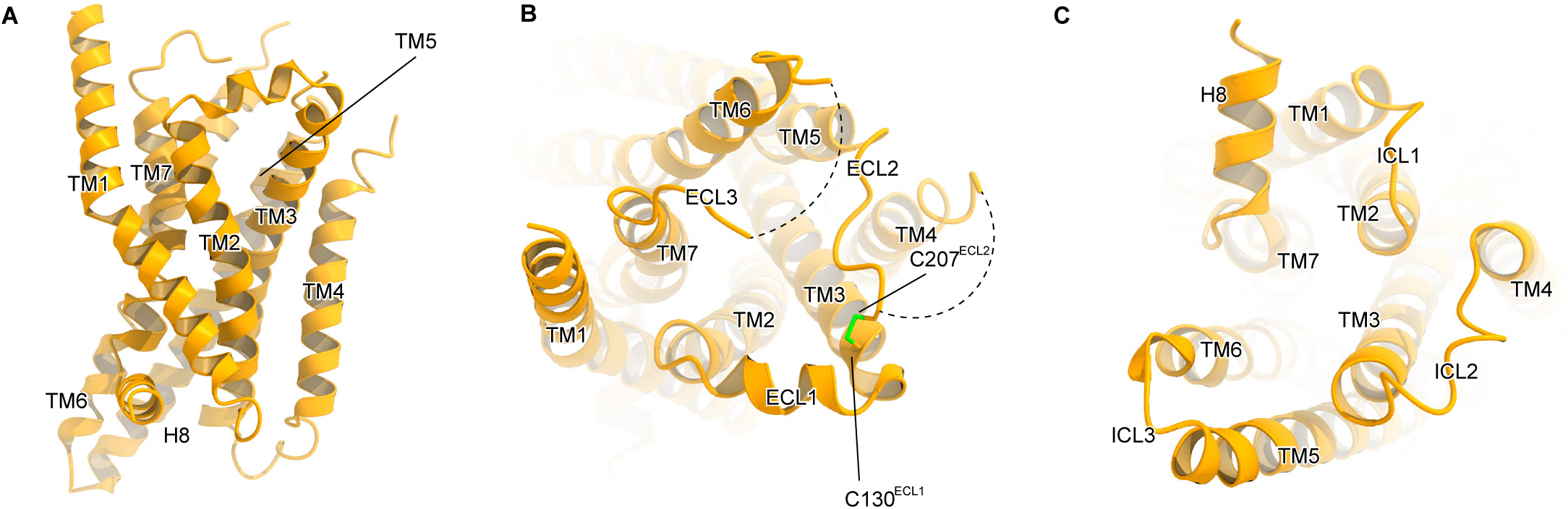
Receptor structure. **A–C** Overall structure of the receptor, viewed from the membrane plane (**A**), intracellular side (**B**), and extracellular side (**C**).

### Extracellular pockets

The interaction network between ECL1–3 covers the extracellular side of the receptor. As described above, ECL2 is anchored to TM3 via the disulfide bond, and ECL3 extends between ECL1 and ECL2, with R299^ECL3^ located within the receptor cavity (Figure 3A, B). R299^ECL3^ forms an electrostatic interaction with D210^ECL2^. These interactions among the ECLs create three extracellular pockets (pockets A–C) (Figure 3A). Pocket B consists of ECL1, TM1, and TM7 (Figure 3C), while pocket C is formed by TM4, TM5, and ECL1 (Figure 3D). These two pockets are small, superficial, and hydrophobic and appear unsuitable for bicarbonate binding. Pocket A consists of ECL1–3 and TM2–7 and is the largest among the three pockets. It is connected to the inside of the receptor, and deep enough to reach W272^6.48^ (Figure 3B). Pocket A contains numerous hydrophilic residues favorable for bicarbonate binding (Figure 3E), suggesting that pocket A is a good candidate for the bicarbonate binding site.

**Figure 3.**
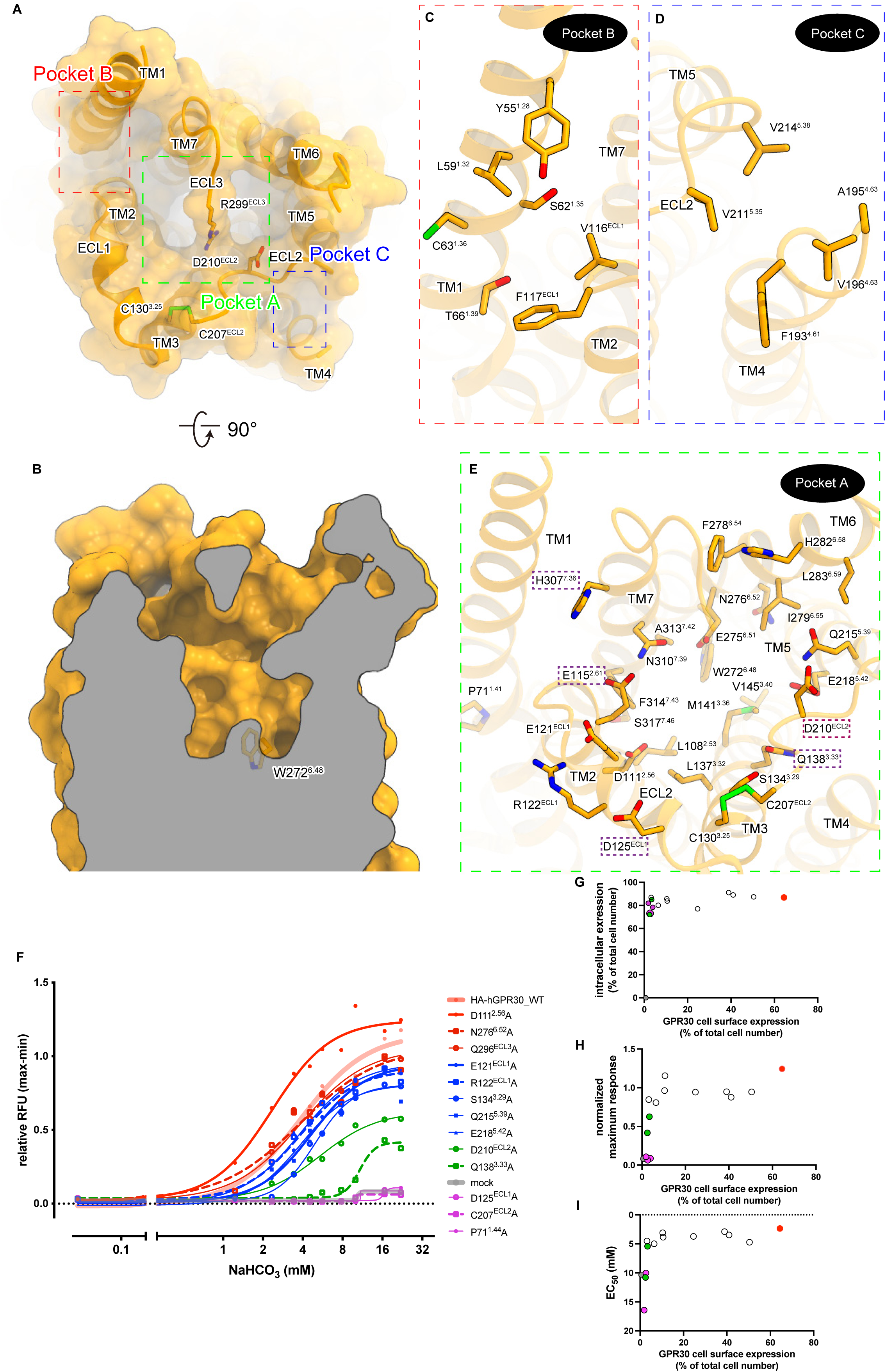
Architecture of the extracellular pocket. **A** Molecular surface of the extracellular side. **B** Cross section of the pocket. **C**–**E** Residues facing pocket B (**C**), pocket C (**D**), and pocket A (**E**). In panel (**E**), only residues with reduced bicarbonate responses are highlighted. **F** Calcium assay using Stable HEK293 cell lines expressing the N-terminal HA-tagged wild-type (WT) and mutant GPR30. The mutants D111^2.56^A, N276^6.52^A, Q296^ECL3^A; E121^ECL1^A, R122^ECL1^A, S134^3.29^A, Q215^5.39^A, E218^5.42^A; D210^ECL2^A, Q138^3.33^A; and D125^ECL1^A, C207^ECL2^A, P71^1.44^A are highlighted in red, blue, green, and purple, respectively. The cells were stimulated by the indicated concentrations of NaHCO_3_ and 50 µM ATP at the timepoint of t = 20 sec. The Y axis indicates the difference between the maximum and minimum fluorescent values during 15 to 60 sec, normalized by those with ATP stimulation. Nonlinear regression (four-parameter) was used for curve-fitting. **G–H** X-Y plot of parameters calculated by FACS analysis and Ca assay. D111^2.56^A; D210^ECL2^A, Q138^3.33^A; D125^ECL1^A, C207^ECL2^A, P71^1.44^A; and mock are colored in red, green, purple, and grey, respectively(**G**). cell surface (x-axis) and total (y-axis) expression of GPR30 (**H**). X-Y plot of cell surface (x-axis) and normalized maximum responses (y-axis) (**I**). X-Y plot of cell surface (x-axis) and EC_50_ (y-axis). EC_50_ was calculated using the data shown in panel (**F**).

However, no obvious density corresponding to bicarbonate is observed in pocket A, along with pockets B and C. In cryo-EM, negative charges are generally difficult to visualize, and the local resolution of the extracellular region in this structure is moderate (Figure 1—figure supplement 2). Thus, even if bicarbonate ions are present, it is possible that they cannot be visualized.

Our previous studies have shown that the mutations of E115^2.61^, Q138^3.33^, and H307^7.36^ abolish the bicarbonate-dependent calcium response of GPR30, suggesting their contribution to bicarbonate binding. To predict the bicarbonate binding site, we performed an exhaustive mutant analysis of the hydrophilic residues in pocket A, which was not done in the previous study^8^. We performed calcium assays using cell lines stably expressing N-terminally HA-tagged GPR30 and its mutants (Figure 3F), whose surface expression levels were analyzed by fluorescence-activated cell sorting (FACS) using an HA-antibody (Figure 3—figure supplement 1-3). We also performed TGFα shedding assays^26^ using HEK cells transiently expressing the receptors (Figure 3—figure supplement 4A-F).

The D125 ^ECL1^A, C207^ECL2^A, and P71^1.44^ A mutations abolished the bicarbonate-dependent calcium responses. The Q138^3.33^A and D210 ^ECL2^A mutations reduced reactivity (Figure 3F). Cell surface expression levels of wild type and mutant GPR30 ranged widely, though the non- and little-responsive mutants, C207^ECL2^A, P71^1.44^A, D125 ^ECL1^A, Q138^3.33^A, and D210 ^ECL2^A showed minimum cell surface expression compared with the others (Figure 3G). We then assessed the correlation between cell surface expression and calcium responses: maximum response (Figure 3H) and EC_50_ (Figure 3I), to examine if the mutants’ reactivity to bicarbonate was attributed to their expression levels on the plasma membrane. Cell surface expression and responsiveness to bicarbonate were not correlated. Q138^3.33^A and D210 ^ECL2^A mutants showed one-third and half maximum responses of GPR30 wild-type, respectively (Figure 3H). The Q138^3.33^A mutant showed a higher EC_50_ value than wild type GPR30, which suggests the involvement in bicarbonate-binding (Figure 3I). D210 ^ECL2^A mutation impairs downstream signaling while keeping its affinity to bicarbonate (Figure 3I). In contrast, the D111^2.56^A mutation enhanced receptor affinity, presenting the half value of wild type EC_50_ (Figure 3.—figure Supplement 4).

Combining these data with previous studies, we mapped the residues that cause decreased bicarbonate responses due to mutations on the current structure (P71^1.44^, E115^2.60^, D125^ECL1^, Q138^3.33^, C207^ECL2^, D210^ECL2^, and H307^7.36^) (Figure 3E). Except for the structure-contributing residues (C207^ECL2^ and P71^1.44^), the essential residues face pocket A, further supporting its role as the bicarbonate binding site. The D210^ECL2^A mutant retained a moderate bicarbonate-induced calcium response (Figure 3F, G) and would contribute to the structural stability of the pocket through its interaction with R299^ECL3^ (Figure 3A). The D125^ECL1^A mutant has lost its activity but is located on the surface (Figure 3E-G) and is unlikely to be a bicarbonate binding site. Given that most bicarbonate binding modes in other structures require recognition by positively charged residues, bicarbonate may bind to H307^7.36^, while E115^2.61^, which is located near H307^7.36^, may play an essential role by coordinating cations for bicarbonate binding, as observed in other structures. These residues are evolutionarily conserved from fish to human, further supporting their importance (Figure 3—figure supplement 5).

### G-protein coupling

The C-terminal helix of Gα_q_ (α5-helix) deeply enters the intracellular cavity of GPR30, resulting in the formation of an active signaling complex. On the C-terminal side of the α5-helix, the backbone carbonyl of Y243^G.H5.23^ (superscript indicates the common Gα numbering [CGN] system^27^) hydrogen bonds with R155^3.50^ in the conserved D^3.49^R^3.50^Y^3.51^ motif^22^ (Figure 4A). In addition, N244^G.H5.24^ forms a hydrogen bond with the peptide backbone of Y324^7.53^. ICL3 faces the middle part of the α5-helix, with three characteristic arginine residues (R248^6.24^, R251^6.27^, and R254^6.30^) surrounding D233^G.H5.13^ (Figure 4B). R248^6.24^ and R254^6.30^ form hydrogen bonds with D233^G.H.13^ and Q237^G.H5.17^, respectively. At ICL2, M163^34.51^ fits into a hydrophobic pocket composed of the α5-helix, the αN-β2 loop, and the β2-β3 loop of the Gα subunit, as in other GPCR-G_q_ complexes^28–32^ (Figure 4C–H). Above it, L159^3.54^ and A162^34.50^ form extensive hydrophobic interactions with L235^G.H5.15^, L236^G.H5.16^, and L240^G.H5.20^ in the α5-helix. These tight interactions with the α5-helix would enable the coupling of GPR30 with G_q_.

**Figure 4.**
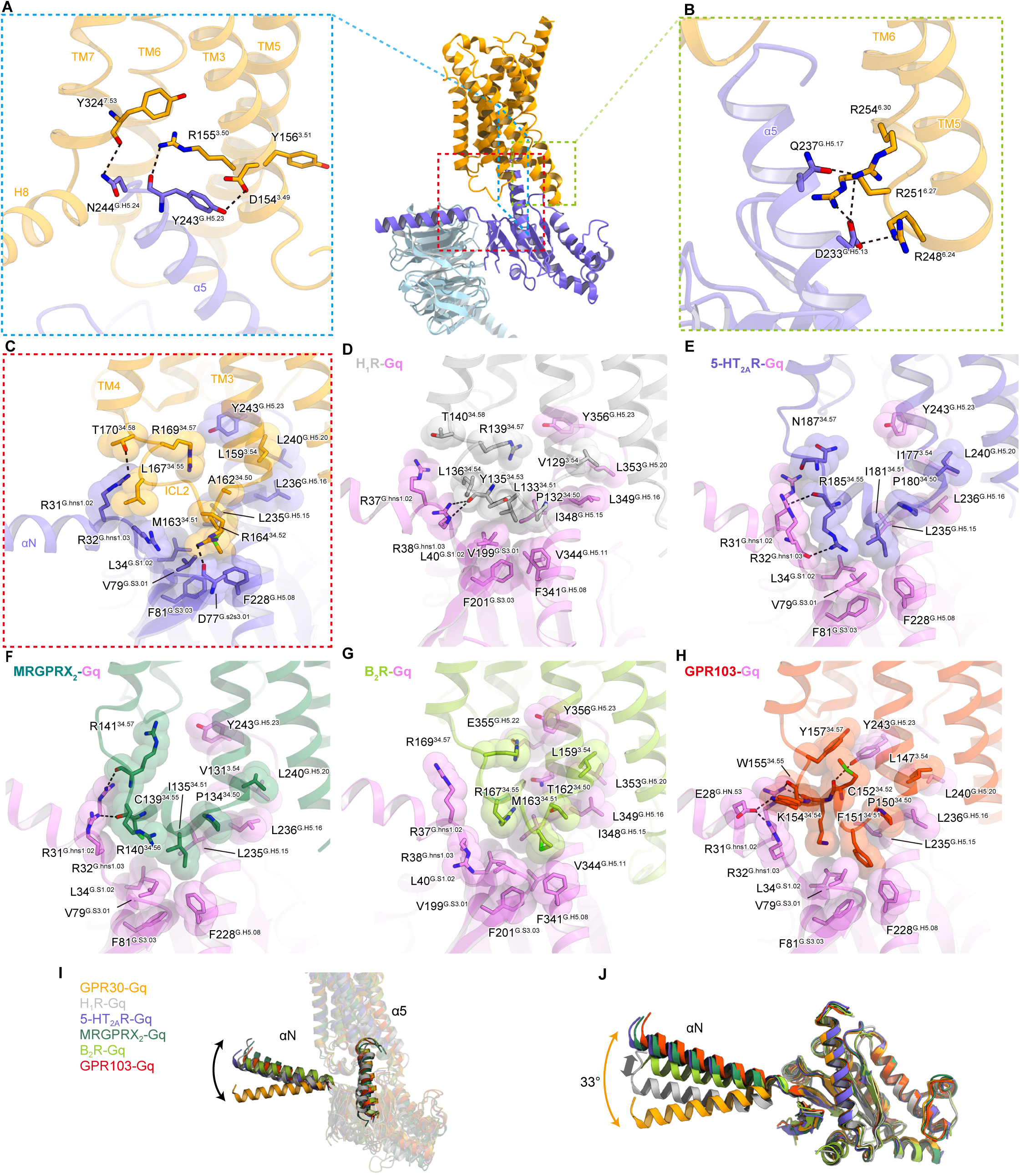
G-protein coupling. **A** Hydrogen-bonding interactions between the C-terminal α5-helix residues and the receptor. **B** Electrostatic and hydrogen-binding interactions between ICL3 and the α5-helix. **C**–**H** Interface between ICL2 and G_q_, with residues involved in hydrophobic interactions represented by CPK models. G-protein-bound GPCRs used in the comparison are as follows: H_1_R-G_q_ (PDB 7DFL, gray), B_2_R-G_q_ (PDB 7F6I, yellow-green), MRGPRX_2_-G_q_ (PDB 7S8L, green), 5-HT_2A_-G_q_ (PDB 6WHA, gray), and GPR103-G_q_ (PDB 8ZH8, red). **I** Comparison of the angles and positions of α5h and αN relative to the receptor. **J** Superimposition of the Gα subunits.

The position of the α5-helix aligns with those in other GPCR-G_q_ complexes, but the position of the αN-helix is rotated away from the receptor (Figure 4I). The superimposition of the Gα subunits indicates that the rotation is not due to the movement of the entire G protein, but rather a 33° bend in the αN-helix (Figure 4J). This is attributed to variations in the interactions of the αN-helix with its receptors. The αN-helix interacts with the intracellular ends of ICL2 and TM4 in receptor-specific modes. In most G_q_-coupled structures^28–32^, ICL2 forms an α-helix and R^34.55^ interacts with R38^G.hns1.03^ at the end of the αN-helix. A typical example is the head-to-toe interaction between R185^34.55^ and R32^G.hns1.03^ in the serotonin 5-HT_2A_ receptor^31^ (Figure 4E). However, in GPR30, ICL2 does not adopt an α-helix and R^34.55^ is replaced by L167^34.55^, which displaces R32^G.hns1.03^ downward through their interaction. Moreover, T170^34.58^ and R169^34.57^ form direct hydrogen bonds with R31^G.hns1.02^ and Y243^G.H5.23^, respectively, which are not observed in other GPCR-G_q_ complexes. We hypothesize that these interactions may underpin the distinct αN rotation: A similar interaction between R38^G.hns1.03^ and L136^34.54^ is also observed in histamine H1 receptor (H_1_R)^28^ (Figure 4D), in which translational rather than rotational motion of αN is observed (Figure 4J).

## Discussion

Our study has elucidated the distinctive structure of GPR30, which is unique among the class A GPCRs. The GPR30 structure is characterized by the kink of TM1 and the presence of the large extracellular pocket covered by ECLs. This pocket is rich in hydrophilic residues and could be a focus for the development of high-affinity drugs targeting GPR30. Our comprehensive mutagenesis study of the binding pocket complemented our previous study and mapped the essential residues for the bicarbonate-induced calcium response. Mutations in the S-S bond between ECL2 and TM3 and the bend in TM1 support their importance in the stability and the shape of the GPR30 structure and pocket. Furthermore, we were able to identify residues in pocket A that are important for the bicarbonate response, although there is room for debate regarding whether they directly contribute to binding or compromise the structure and thereby downstream signaling. Considering the binding sites of negatively charged bicarbonate in other protein structures, it is reasonable to regard the region around H307^7.36^ as a bicarbonate binding site candidate. To clarify the binding mode of bicarbonate, a higher resolution structural analysis and complementary functional assessments are necessary.

The C207^ECL2^A and P71^1.44^ mutation disrupts the disulfide bond between ECL2 and TM3 and the characteristic kink in TM1, respectively. Loss of bicarbonate-dependent activation by these mutations underscores the critical role of the structural integrity of the receptor and provides compelling evidence that the bicarbonate response is mediated through GPR30. Higher EC_50_ value of Q138^3.33^A mutant than that of wild type suggests that the residue might be involved in bicarbonate-binding. D210 ^ECL2^A mutation impairs downstream signaling while keeping its affinity to bicarbonate. Overall, cell surface expression and response to bicarbonate were not correlated as long as only a small portion (∼10%) was expressed on the cell surface. Many possibilities could explain this: GPR30 localization in specific spots on the plasma membrane might limit the response stoichiometry, GPR30 might also work intracellularly to blunt the increased response because of more GPR30 expression on the plasma membrane, redundant GPR30 on the plasma membrane might be broken, or the mutants might be less functional than wild type receptor and need higher expression.

GPR30 exhibits low homology to other GPCRs, with even closely related receptors sharing less than 30% sequence identity. A BLAST analysis revealed that GPR30 shares the highest sequence identity of 28% with the type 2 angiotensin II receptor (AT_2_R) (Figure 5—figure supplement 1). Consequently, we compared the current structure of GPR30 with the ligand-bound AT_2_R^33^. The superimposition of GPR30 and AT_2_R, with a Cα R.M.S.D. of 1.73 □, suggests significant structural divergence between the two (Figure 5A). Notably, there is no structural commonality between GPR30 and AT2R in terms of the orientations of their respective TMs. Specifically, the extracellular half of TM1 in GPR30 extends outward, and is kinked at P71^1.44^ in the center of TM1 (Figure 5B). Near P71^1.44^, Y65^1.38^ and F70^1.43^ are oriented toward the interior of the receptor, forming hydrophobic interactions with L113^2.58^ and L331^7.40^ to stabilize the kink. In the examination of the loops, ECL2 of AT_2_R features a short β-sheet commonly found in peptide-activated class A GPCRs^34–36^, whereas that of GPR30 adopts an elongated conformation that covers the extracellular pocket essential for the bicarbonate response (Figure 5A, B). These comparisons underscore the unique structural characteristics of GPR30.

**Figure 5.**
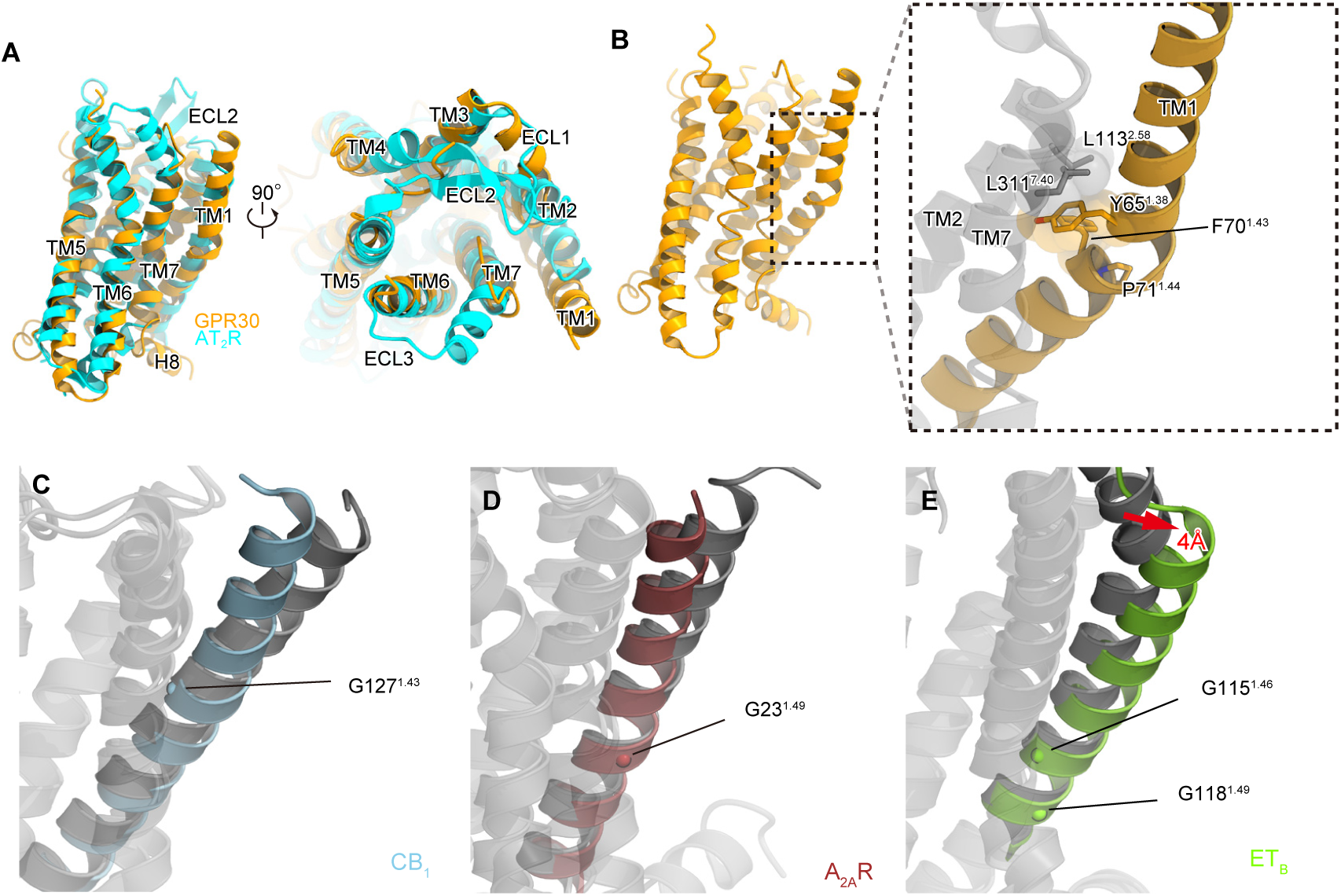
Structural comparison with related GPCRs. **A** Structural comparison of GPR30 and AT_2_R (PDB 5UNF). **B** Interactions around P71^1.44^ in TM1 of GPR30. **C–F** Conformational changes of TM1 upon agonist binding in CB1 (**C**), A_2A_R (**D**), and ET_B_ (**E**). The agonist-bound states are colored with the respective colors, while the inactive states are colored gray. The PDB codes used in this figure are CB1-active (PDB 5XRA), CB_1_-inactive (PDB 5TGZ), A_2A_R-active (PDB 6GDG), A_2A_R-inactive (PDB 3EML), ET_B_-active (PDB 8IY5), and ET_B_-inactive (PDB 5X93).

In most GPCRs, TM1 adopts a straight α-helix and is positioned away from the orthosteric pocket formed by TM2–7. However, TM1 undergoes an inward movement due to allosteric conformational changes during agonist binding. The cannabinoid receptor CB_1_ and adenosine A2A receptor (A_2A_R) exemplify this phenomenon, with TM1 kinked at G127^1.43^ and G23^1.49^ folding inward upon ligand binding^37–40^ (Figure 5C, D). By contrast, TM1 moves 4 Å outward in the endothelin ET_B_ receptor^41,42^ (Figure 5E). In GPR30, TM1 is kinked at P71^1.44^ (Figure 5B), indicating that this kink exists regardless of the receptor activation state. Indeed, the P71^1.44^A mutation, which eliminates the kink in TM1, completely abolished the bicarbonate response (Figure 3F, G). Moreover, P71^1.44^ is evolutionarily conserved (Figure 3—figure supplement 5). These findings suggest the critical role of the TM1 kink in both the putative conformational changes associated with ligand binding and the functional integrity of GPR30.

During the preparation of the manuscript, the structures of apo-GPR30-G_q_ (PDB 8XOG) and the exogenous ligand Lys05-bound GPR30-G_q_ (PDB 8XOF) were reported^43^. We compared our structure of GPR30 in the presence of bicarbonate with these structures. In the extracellular region, the position of TM5 in GPR30 in the presence of bicarbonate is similar to that in apo-GPR30. In contrast, the position of TM6 is shifted outward relative to that of apo-GPR30, resembling the conformation observed in Lys05-bound GPR30 (Figure 6A, B). Additionally, the position of ECL1 is also shifted outward compared to that of apo-GPR30 (Figure 6B). In the GPR30 structure in the presence of bicarbonate, ECL2 was modeled, suggesting differences in structural flexibility. These findings indicate that the structure of GPR30 in the presence of bicarbonate is different from both the apo structure and the Lys05-bound structure, demonstrating that the structure and the flexibility of the extracellular domain of GPR30 change depending on the type of ligand. Furthermore, focusing on the interaction with G_q_, the αN helix of G_q_ is not rotated in the structure bound to Lys05, in contrast to the characteristic bending of the αN helix in our structure (Figure 6C, D). Although it is necessary to consider variations in experimental conditions, such as salt concentration, the differences in the G_q_ binding modes suggest that the downstream signals may change in a ligand-dependent manner.

**Figure 6.**
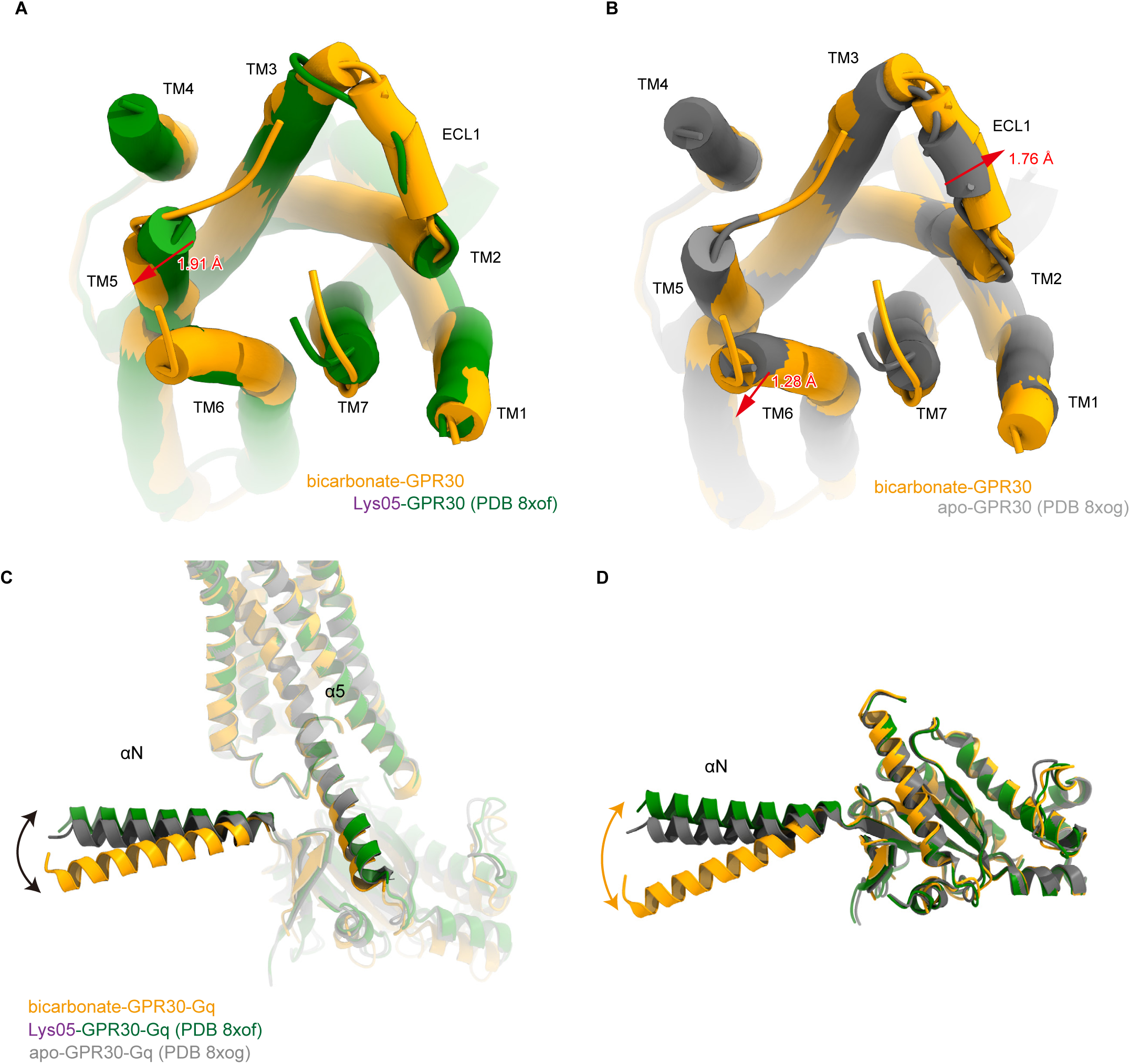
Structural comparison with GPR30 bound to Lys05. **A**, **B** Superimposition of bicarbonate-GPR30, Lys05-binding GPR30 (PDB 8XOF) (**A**), and apo-GPR30 (PDB 8XOG) (**B**) structures, viewed from the extracellular side. **C** Comparison of the angles and positions of α5h and αN relative to the receptor. **D** Superimposition of the Gα subunits.

In our work, we used relatively high (mM) concentrations of bicarbonate to activate GPR30. In parallel with its low potency, the physiological concentrations of bicarbonate are 22-26 mM in the extracellular fluid, including interstitial fluid and blood, and 10-12 mM in the cytoplasm. Therefore, GPR30 is activated by the physiological concentrations of bicarbonate in tissues. The bicarbonate concentrations are altered in various physiological and pathological conditions – metabolic acidosis in chronic kidney disease causes a drop to 2-3 mM, and metabolic alkalosis induced by severe emesis increases bicarbonate concentrations to over 30 mM. Thus, our work clearly shows that GPR30 is activated by physiological concentrations of bicarbonate, present both intracellularly and extracellularly, and that GPR30 can be deactivated or reactivated under various pathophysiological conditions. Thus, we have identified GPR30 as a bicarbonate receptor.

---

**Figure 1.—figure supplement 1.** Sample purification. **A** Schematic representation of the fusion-G system. **B** Fluorescence-detection size-exclusion chromatography (FSEC) analysis of complex formation by GPR30. The trace from solubilized cells expressing only GPR30 is orange, and that from cells co- expressing GPR30 and G-protein is blue. **C** Size-exclusion chromatography of the GPR30-G-protein complex on a Superose 6 Increase column. The fraction shaded in blue was collected. **D** SDS-PAGE gel of samples after size-exclusion chromatography, stained with Coomassie Brilliant Blue.

**Figure 1.—figure supplement 2.** Cryo-EM analysis. Flow chart of the cryo-EM data processing for the GPR30-G_q_ complex, including particle projection selection, classification, and 3D density map reconstruction. The 3D density map was refined with a mask on the receptor. Details are provided in the Methods section.

**Figure 2.—figure supplement 1.** ECL2 density. **A-C** top view of density map around disulfide bond between ECL2 and TM3 at counter levels 8.5 (**A**), 7.25 (**B**), and 6 (**C**). **D-F** side view of density map around disulfide bond between ECL2 and TM3 at counter levels 8.5 (**D**), 7.25 (**E**), and 6 (**F**).

**Figure 2.—figure supplement 2.** Structural comparison with AlphaFold-predicted structure. **A–C** Superimposition of the cryo-EM (orange) and AF3 (khaki) structures. (**A**) Overall view of the receptor, (**B**) focused on the intracellular side, and (**C**) focused on the extracellular side.

**Figure 3.—figure supplement 1. Cell surface expression of stable wild-type and GPR30 mutants.** Flow cytometric-based assay to analyze WT and mutant GPR30 cell surface expression. Cell surface expression, defined as the HA-positive subset (red) compared with isotype control staining (blue), is indicated on each histogram.

**Figure 3.—Figure Supplement 2. Flow cytometric gating criteria to analyze GPR30 cell surface expression. A**, **B** Gating used in the analysis. **C**, **D** Cell surface expression is defined as the HA-positive subset (red) compared with isotype control staining (blue). **(C)** Mock cells, (**D**) WT GPR30-expressing cells.

**Figure 3.—Figure Supplement 3. Flow cytometric-based assay to analyze the whole-cell expression of WT GPR30 and its mutants. A** Gating used in the analysis. **B** Whole-cell expression is defined as the HA-positive subset (red) compared with isotype control staining (blue).

**Figure 3.—Figure Supplement 4. Cell surface expression and bicarbonate-induced activation of HEK293 cells transiently expressing GPR30 mutants. A, B** TGFα shedding assay using HEK293 cells transfected with HA-tagged hGPR30. The mutants D111^2.56^A; D125^ECL1^A, C207^ECL2^A, P71^1.44^A, and H307^7.36^A; and S134^3.29^A, D210^ECL2^A, and Q215^5.39^A are highlighted in red, blue, and purple, respectively. **C** Whole-cell expression of HA-tagged mutants, analyzed by western blotting with 20 μg of whole-cell lysate per lane. **D** Cell surface expression of HA-tagged mutants, using cell surface biotinylation and avidin immunoprecipitation, analyzed by western blotting with 1.5 μg of cell surface protein per lane. **E, F** Quantitative analysis of HA expression in **C** and **D**.

Statistical analysis: ^$^ *p* = 0.01, ^§^ *p* < 0.001 compared to HA-hGPR30 cells stimulated with vehicle, ^#^ *p* < 0.0001 compared to HA-hGPR30 cells stimulated with 11 mM NaHCO_3_, using two-tailed unpaired t-test with Bonferroni’s correction after two-way ANOVA. Data are presented as mean values (**A**) and mean values ± SEM (**B**).

**Figure 3.—figure supplement 5. Conservation of the GPR30 homologs.** Amino acid alignment of representative GPR30 homologs.

**Figure 5.—figure supplement 1. Sequence alignment of GPR30 and AT_2_R.**

## Data Availability

The cryo-EM density map and atomic coordinates for the GPR30-G_q_ complex have been deposited in the Electron Microscopy Data Bank and the PDB, under the accession codes: EMD-66632 [https://www.ebi.ac.uk/emdb/entry/EMD-66632] (GPR30-G_q_), EMD-66629 [https://www.ebi.ac.uk/emdb/entry/EMD-66629] (GPR30-G_q_, consensus map), EMD-66630 [https://www.ebi.ac.uk/emdb/entry/EMD-66630] (GPR30-G_q_, G-protein focused), EMD-66631 [https://www.ebi.ac.uk/emdb/entry/EMD-66631] (GPR30-G_q_, receptor focused), and PDB 9X74 [http://doi.org/10.2210/pdb9X74/pdb]. Source data are provided with this paper. All other data are available from the corresponding authors upon reasonable request.

## Acknowledgements

We thank K. Ogomori and C. Harada for technical assistance. This work was supported by grants from the JSPS KAKENHI, grant numbers 21H05037 (O.N.), 25K02397 (W.S.), 24KJ0906 (H.A.), 23K06393, 24KK0146, and 25H01336 (T.Y); the Japan Agency for Medical Research and Development (AMED), grant numbers JP233fa627001 (O.N.), 25ak0101252h0001 (W.S,), and JP20gm6210026 (A.J.-W.); the JST FOREST Program (Grant number JPMJFR220S to A.J.-W.); the Takeda Science Foundation (W.S. and A.J.-W.); the Mochida Memorial Foundation for Medical and Pharmaceutical Research (W.S.); Dojindo Laboratories’ Foundation for Life Science (W.S.); the Nakatani Foundation (W.S.); the Mitsubishi Foundation (W.S.); the Daiichi Sankyo Foundation of Life Science (W.S.); the Takahashi Industrial and Economic Research Foundation (W.S.); G-7 Scholarship Foundation (W.S.); and the Platform Project for Supporting Drug Discovery and Life Science Research (Basis for Supporting Innovative Drug Discovery and Life Science Research (BINDS)) from AMED, grant numbers JP23ma121002 and JP23ama121012.

## Author contributions

S.K. performed all the experiments involved in the structural determination, assisted by H.A. and H.S.O. W.S. designed the project and initially tried the expression of GPR30. A.J.-W. and T.Y. performed and oversaw the mutagenesis study. The manuscript was mainly prepared by S.K., H.A., and W.S., with assistance from O.N.

## Competing interests

The authors declare no competing interests.

## Methods

### Expression and purification of scFv16 and Nb35

The gene encoding scFv16 was synthesized (GeneArt) and subcloned into a modified pFastBac vector^44^, with the resulting construct encoding the GP67 secretion signal sequence at the N terminus, and a His_8_ tag followed by a TEV cleavage site at the C terminus. The His_8_-tagged scFv16 was expressed and secreted by Sf9 insect cells, as previously reported^45^. The Sf9 cells were removed by centrifugation at 5,000g for 10□min, and the secreta-containing supernatant was combined with 5□mM CaCl_2_, 1 mM NiCl_2_, 20□mM HEPES (pH□8.0), and 150□mM NaCl. The supernatant was mixed with Ni Superflow resin (GE Healthcare Life Sciences) and stirred for 1 h at 4□□. The collected resin was washed with buffer containing 20□mM HEPES (pH□8.0), 500□mM NaCl and 20□mM imidazole, and further washed with 10□column volumes of buffer containing 20□mM HEPES (pH□8.0), 500□mM NaCl and 20□mM imidazole. Next, the protein was eluted with 20□mM Tris (pH□8.0), 500□mM NaCl and 400□mM imidazole. The eluted fraction was concentrated and loaded onto a Superdex200 10/300 Increase size-exclusion column, equilibrated in buffer containing 20□mM Tris (pH□8.0) and 150□mM NaCl. Peak fractions were pooled, concentrated to 5□mg/ml using a centrifugal filter device (Millipore 10□kDa□MW cutoff), and frozen in liquid nitrogen.

Nb35 was prepared as previously reported^46,47^. In brief, Nb35 was expressed in the periplasm of *E. coli*. The harvested cells were disrupted by sonication. Nb35 was purified by nickel affinity chromatography, followed by gel-filtration chromatography, and frozen in liquid nitrogen.

### Constructs for expression of GPR30 and G_q_

The human GPR30 gene (UniProtKB, Q99527) was subcloned into a modified pFastBac vector^20^, with an N-terminal haemagglutinin signal peptide followed by the Flag-tag epitope (DYKDDDDK) and the LgBiT fused to its C-terminus followed by a 3 C protease site and EGFP-His_8_ tag. A 15 amino sequence of GGSGGGGSGGSSSGG was inserted into both the N-terminal and C-terminal sides of LgBiT. Rat Gβ_1_ and bovine Gγ_2_ were subcloned into the pFastBac Dual vector. In detail, rat Gβ_1_ was cloned with a C-terminal HiBiT connected with a 15 amino sequence of GGSGGGGSGGSSSGG. Moreover, mini-G_sqi_ was subcloned into the C-terminus of the bovine Gγ_2_ with a nine amino sequence GSAGSAGSA linker. The resulting pFastBac dual vector can express the mini-G_sqi_ trimer.

### Expression and purification of the human GPR30 – G_q_ complex

The recombinant baculovirus was prepared using the Bac-to-Bac baculovirus expression system (Thermo Fisher Scientific). For expression, 2 L of Sf9 cells at a density of 3 × 10^6^ cells/mL were co-infected with baculovirus encoding GPR30 and mini-G_sqi_ trimer at the ratio of 1:1 and the cells were incubated at 30 □. After 48 hours, the collected cells were resuspended and dounce-homogenized in 20 mM Tris-HCl, pH□8.0 (4 □), 200□mM NaCl, 10% Glycerol, 200 mM NaHCO_3_, 25 mU/ml apyrase. The crude membrane fraction was collected by ultracentrifugation at 180,000*g* for 1□h and solubilized in buffer, containing 20□mM Tris-HCl, pH□8.0, 150□mM NaCl, 1% n-dodecyl-beta-D-maltopyranoside (DDM) (Calbiochem), 0.2 % cholesteryl hemisuccinate (CHS) (Merck), 10% glycerol, 200 mM NaHCO_3_, and 25 mU/ml Apyrase, for 1 h at 4□□. The supernatant was separated from the insoluble material by ultracentrifugation at 180,000*g* for 20□min and incubated with 5 mL of M2 anti-flag affinity resin (Sigma) for 1 h at 4 □. The resin was washed with 20 column volumes of buffer containing 20□mM Tris-HCl, pH□8.0 500□mM NaCl, 10% Glycerol, 0.05% Lauryl Maltose Neopentyl Glycol (LMNG) (Antrace), 0.005% CHS and 200 mM NaHCO_3_. The complex was eluted in buffer containing 20□mM Tris-HCl, pH□8.0, 150□mM NaCl, 10% Glycerol, 0.01% LMNG, 0.001% CHS, 200 mM NaHCO_3_ and 0.2 mg/mL Flag peptide. The eluate was incubated with the Nb35 and scFv16 at 4 □. The complex was concentrated and purified by size exclusion chromatography on a Superose 6 increase (GE) column in 20□mM Tris-HCl, pH8.0, 150□mM NaCl, 0.01% LMNG, 0.001% CHS and 200 mM NaHCO_3_. Peak fractions were concentrated to 4.72 mg/ml for electron microscopy studies.

### Sample vitrification and cryo-EM data acquisition

The purified complex was applied onto a freshly glow-discharged Quantifoil Au grid (R1.2/1.3, 300 mesh), and plunge-frozen in liquid ethane by using a Vitrobot Mark IV. Data collections were performed on a 300kV Titan Krios G3i microscope (Thermo Fisher Scientific) equipped with a BioQuantum K3 imaging filter and a K3 direct electron detector (Gatan).

A total of 9824 movies were acquired with a calibrated pixel size of 0.83□Å□pix^-1^ and with a defocus range of -0.8 to -1.6□μm, using EPU. Each movie was acquired for 2.6□s and split into 48 frames, resulting in an accumulated exposure of about 50.660□electrons per Å^2^.

### Image processing

All acquired dose-fractionated movies were imported into CryoSPARC v4.4^48^ and subjected to beam-induced motion correction. The contrast transfer function (CTF) parameters were estimated and a total of 10,148,422 particles were extracted. The particles were subjected to 2D classifications, Ab-initio reconstruction and several rounds of hetero refinement and Non-uniform refinement. Next, the particles were subjected to 3D classification with a mask on the receptor. Then the particle sets were subjected to Reference Based Motion Correction. Motion-corrected 522,404 particles were subjected to Non-uniform refinement, yielding a map with a global nominal resolution of 3.15 Å, with the gold standard Fourier Shell Correlation (FSC=0.143) criteria^49^. Moreover, the 3D model was refined with a mask on the receptor. As a result, the receptor has a higher resolution with a nominal resolution of 3.18 Å. The overall and receptor focused maps were combined by phenix^50^. The processing strategy is described in Supplementary Figure 2.

### Model building and refinement

The density map was sharpened by phenix.auto_sharpen^51^ and the quality of the density map was sufficient to build a model manually in COOT^52,53^. The model building was facilitated by the Alphafold-predicted structure. We manually fitted GPR30, the G_q_ heterotrimer and scFv16 into the map. We then manually readjusted the model into using COOT and refined it using phenix.real_space_refine^50,54^ (v.1.19) with the secondary-structure restraints using phenix secondary_structure_restraints.

### Vector construction and transfection

Human *Gpr30* cDNA was obtained from human hepatoblastoma-derived HepG2 cells. The coding sequences of human *Gpr30* were inserted into the multi-cloning site of the plasmid vector pCXN2, which was generated in our laboratory via modification of pCAGGS, between the KpnI and EcoRI sites. The N- and C-terminal HA-tagged *Gpr30* was amplified using a reverse primer containing the HA sequenceOne amino acid mutation of human GPR30 was generated as follows: the targeted amino acid was changed to alanine (GCC) using a two-step PCR method with the QuikChange® Primer Design Program by Agilent (https://www.agilent.com/store/primerDesignProgram.jsp), and the coding sequence with each mutation was inserted into the multi-cloning site of pCXN2 between the KpnI and EcoRI sites. The mutations generated were D111A, E121A, R122A, D125A, S134A, Q138A, D210A, Q215A, E218A, N276A, Q296A, C207A, P71A, and H307A.

### Cell line sources and transfection

HEK293A cells (female origin; Thermo Fisher Scientific) were maintained at 37 □ and 5% CO_2_. These vectors were transfected using the lipofection method (Lipofectamine™ 2000 Transfection Reagent, 11668019, Invitrogen). Stable cell lines were established through antibiotic drug selection. The HEK293A cells were transfected with the N-terminal HA-tagged wild-type and mutant *Gpr30* and maintained in the culture medium containing 1 mg/ml of G418. After 2-3 weeks of antibiotic drug selection, stable expression of exogenous GPR30 was confirmed by three independent flow cytometric analyses.

### TGFα shedding assay

The TGFα shedding assay was performed according to a previously published protocol^26^. The AP-TGFα expression vector was provided by Dr. Inoue and Dr. Aoki, Tohoku University. HEK293 cells were seeded in 12-well plates at a density of 1×10^5^ cells/well and cultured for 24 h. At 70% confluency, a mixture of plasmid vectors containing GPCR (see ‘***Vector construction and transfection****’* section) and AP-TGFα was transfected into the cells using Lipofectamine 2000 transfection reagent. After another 24 h of incubation, the cells were detached with 0.05% trypsin/EDTA (32777-44, Nacalai Tesque), suspended in Hanks’ balanced salt solution, and seeded in 96-well plates. The cells were stimulated for 1 h with 1–44×10^-3^ M (final concentration) of NaHCO_3_ at 37 □, under 0.03% CO_2_. Conditioned media (CM) was transferred to another plate, and 80 µl of alkaline phosphatase (AP) solution (40 mM Tris-HCl, pH 9.5, 40 mM NaCl, 10 mM MgCl_2_, 10 mM p-nitrophenylphosphate disodium salt hexahydrate) was added to both plates, which were then incubated at 37 □. The optical density at 405 nm (OD405) was measured using a microplate reader (iMark, Bio-Rad) at 5 min, 30 min, 1 h, and 2 h, depending on the reaction rate. The percentage of shed AP-TGF was calculated using the following equations:

AP activity = ΔOD405 (1 – 0 h)

% CM (conditioned media) = AP_CM_ / (AP_CM_ + AP_Cell_)

### Calcium assay

Stable HEK293 cells (2×10^4^ cells/well) were seeded into a black wall and clear bottom 96-well plate 24 h before the assay. Cells at 90–100% confluency were incubated with HEPES buffer (1× HBSS, 2.5 mM probenecid, 25 mM HEPES, pH 7.4) containing 10 µM Fluo-8 AM (21080, AAT-Bio) for 60 min at 37□ and 5% CO_2_. The cells were washed twice with HEPES buffer. The cells were stimulated with indicated concentrations of NaHCO_3_ and 50 µM ATP at 37 □ and 0.03% CO_2_. The fluorescence intensity was analysed using FlexStation 3 (Molecular Devices, Ex/Em = 490/525 nm). The difference between the maximum and minimum fluorescent values during 15 to 60 sec was normalized by those with ATP stimulation. Nonlinear regression (four-parameter) was used for curve-fitting, and EC_50_ and Top (maximum response) were calculated.

### Flow cytometry

HEK293A cells stably expressing the N-terminal HA-tagged wild-type and mutant GPR30 were incubated with the calcium-free Minimum Essential Medium (S-MEM, #11380037, Gibco™) for 30 min at 37 □ and 5.0% CO_2_ before the assay. The cells were blocked with PBS containing 2% goat serum for 10 min at 4□, incubated with the biotin-conjugated anti-HA antibody (2.5 µg/ml, #12158167001, Roche) for 60 min at 4□, and then stained with Alexa Fluor 488-conjugated anti-rat IgG (5 µg/ml, #A-11006, Thermo Fisher Scientific) for 40 min at 4□. LSRFortessa (BD Biosciences) was used for analysis. For intracellular staining, the cells were fixed and stained using eBioscience™ Fixable Viability Dye eFluor 780 (#65-0865-14, Thermo Fisher Scientific) and eBioscience™ Fox3/Transcription Factor Staining Buffer Set (#00-5523-00, Thermo Fisher Scientific).

### Analysis of cell surface expression of GPR30

The cell surface expression of wild-type and mutant hGPR30-HA was analysed via cell surface protein isolation using a Cell Surface Protein Isolation Kit (#89881, Thermo Scientific™) followed by western blotting. Briefly, HEK293 cells transiently expressing wild-type or mutant hGPR30-HA were biotinylated for 30 min at 4 □. Then, the cells were harvested, and biotinylated proteins were isolated with avidin binding.

### Western blotting

Cell lysates were separated using sodium dodecyl sulfate-polyacrylamide gel electrophoresis under reducing conditions and transferred to polyvinylidene difluoride membranes (Immobilon P, IPVH00010, Millipore). Primary antibodies used were anti-HA High Affinity, 1:1000 dilution, 11867423001, Roche; Na-K-ATPase, 1:1000 dilution, #3010, Cell Signaling Technology; β-Actin (AC-15), 1:1000 dilution, sc-69879, Santa Cruz Biotechnology. Secondary antibodies used were anti-rabbit IgG, HRP-linked Antibody, 1:5000 dilution, #7074, Cell Signaling Technology; anti-mouse IgG, HRP-linked Antibody, 1:5000 dilution, #7076, Cell Signaling Technology; anti-rat IgG, HRP-Linked Whole Ab Goat, 1:5000 dilution, NA935, Cytiva. The membranes were probed at 4 □ overnight with the primary antibodies. The membranes were subsequently incubated with the corresponding secondary antibodies. The signals were detected with ECL Prime (RPN2236, Cytiva) or ImmunoStar LD (296-69901, FUJIFILM Wako Pure Chemical Corporation) using a chemiluminescence imaging system (CFusion FX7, Vilber).

## Notes

### Competing Interest Statement

The authors have declared no competing interest.

### Summary of Updates

Figure 2,3 revised; Supplemental files updated.

