## Supplemental figure for "Cryo-EM structure of the bicarbonate receptor GPR30"

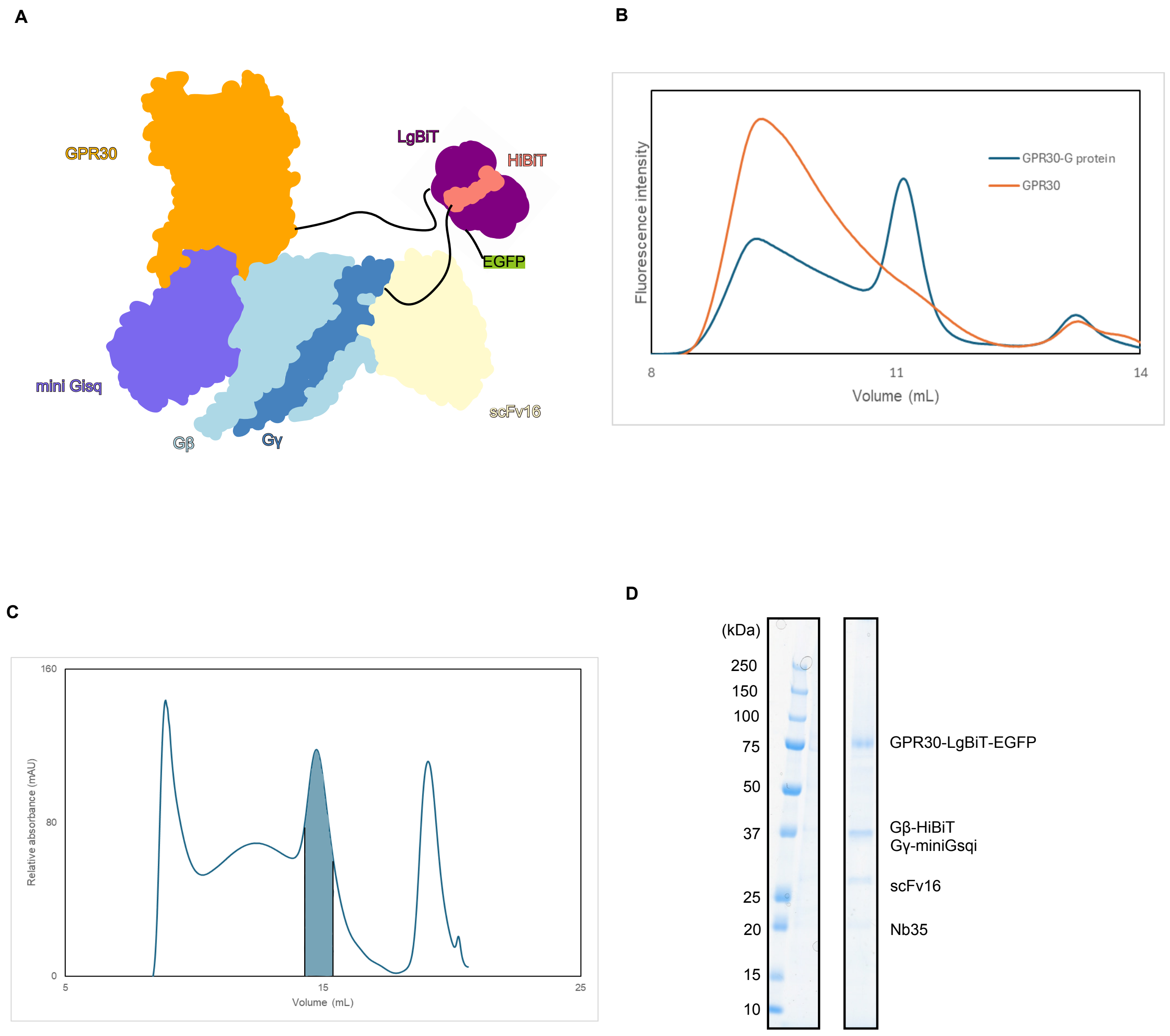


**Figure 1.—figure supplement 1. Sample purification.**
**A** Schematic representation of the fusion-G system. **B** Fluorescence-detection size-exclusion chromatography (FSEC) analysis of complex formation by GPR30. The trace from solubilized cells expressing only GPR30 is orange, and that from cells co-expressing GPR30 and G-protein is blue. **C** Size-exclusion chromatography of the GPR30-G-protein complex on a Superose 6 Increase column. The fraction shaded in blue was collected. **D** SDS-PAGE gel of samples after size-exclusion chromatography, stained with Coomassie Brilliant Blue.

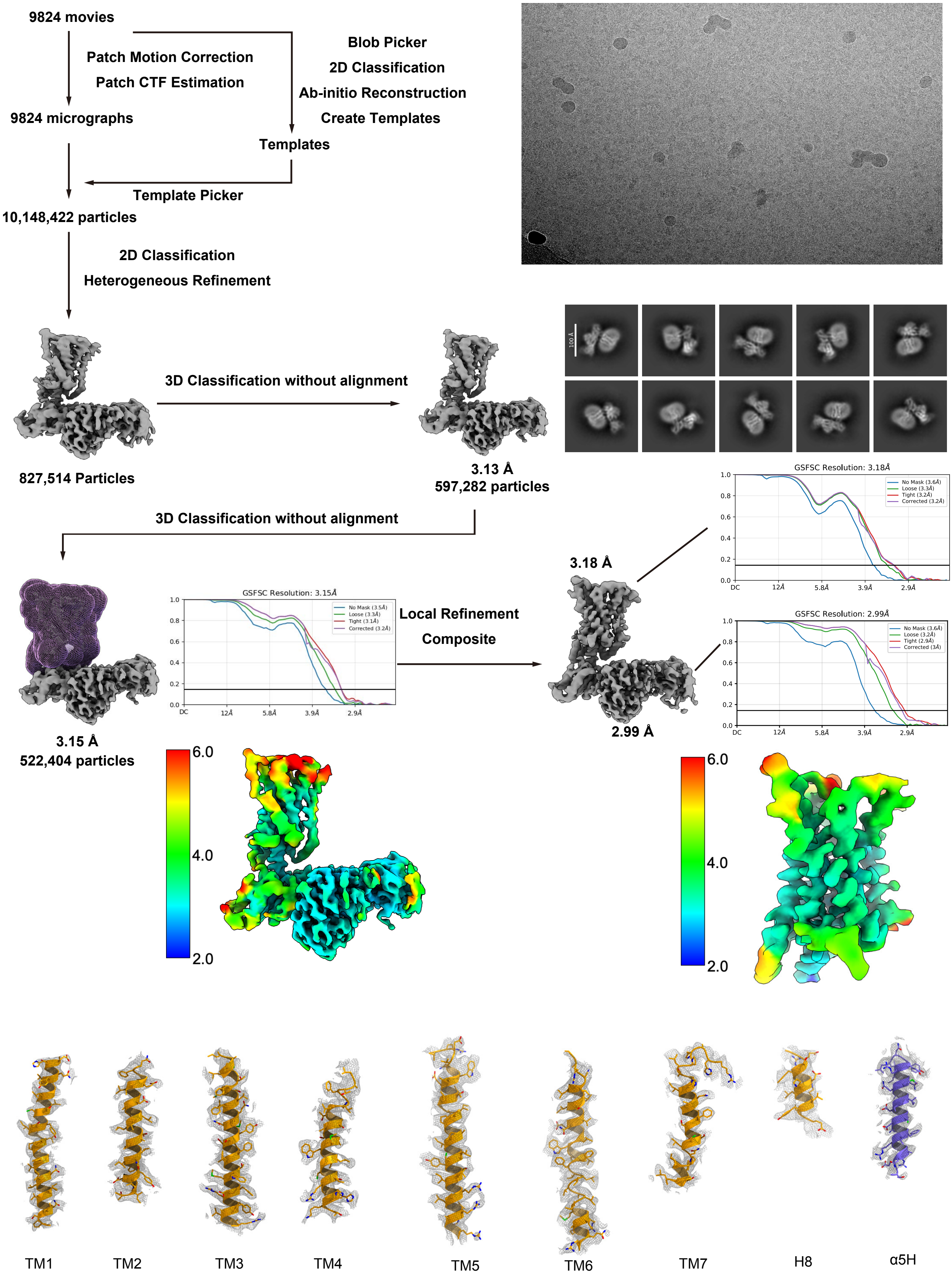


**Figure 1. —figure supplement 2. Cryo-EM analysis.**
Flow chart of the cryo-EM data processing for the GPR30-$G_q$ complex,
including particle projection selection, classification, and 3D density map reconstruction.
The 3D density map was refined with a mask on the receptor.
Details are provided in the Method section.

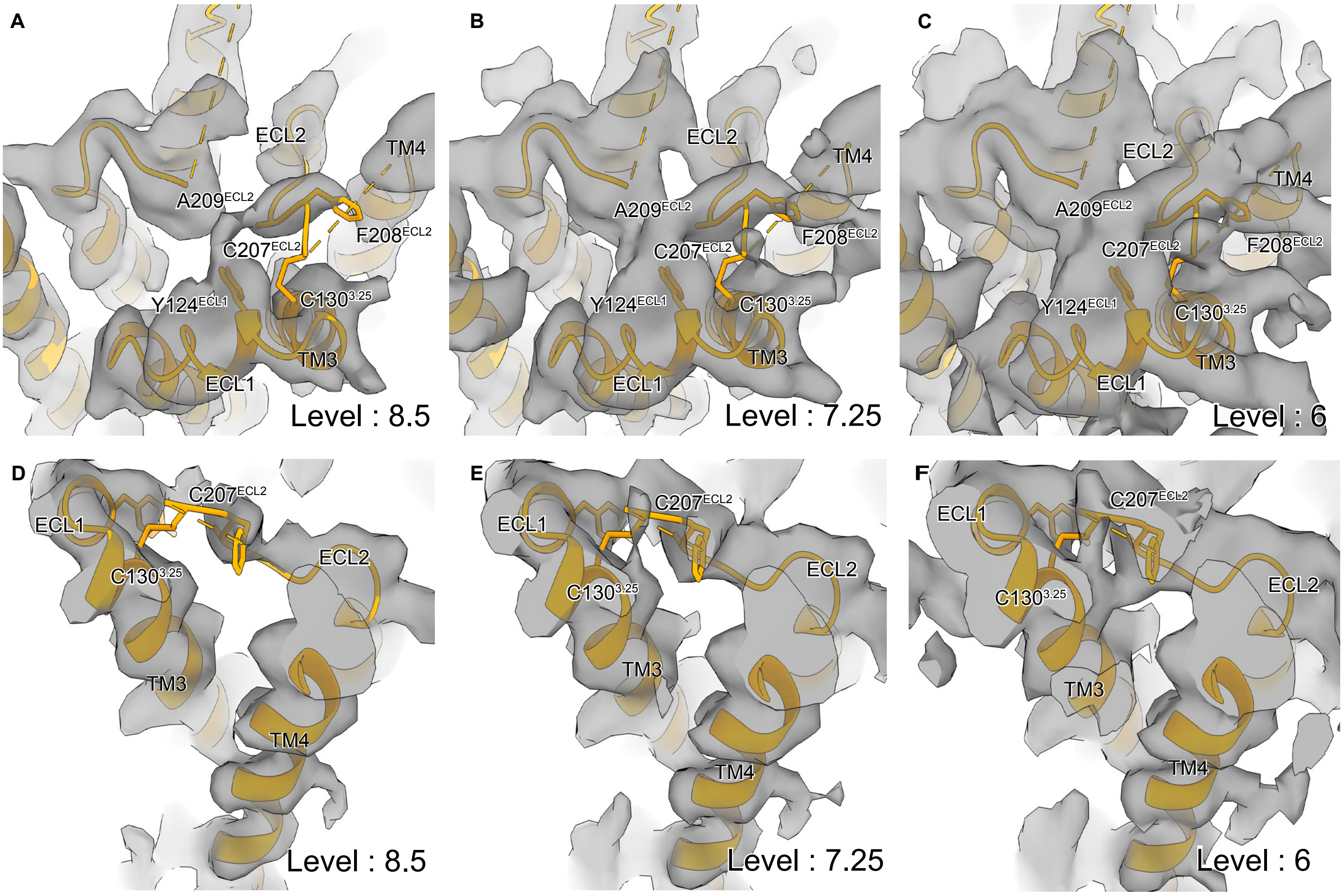


**Figure 2.—figure supplement 1. ECL2 dencity.**
**A-C** top view of density map around disulfide bond between ECL2 and TM3 at counter levels 8.5 (**A**), 7.25 (**B**), and 6 (**C**)**. D-F** side view of density map around disulfide bond between ECL2 and TM3 at counter levels 8.5 (**D**), 7.25 (**E**), and 6 (**F**).

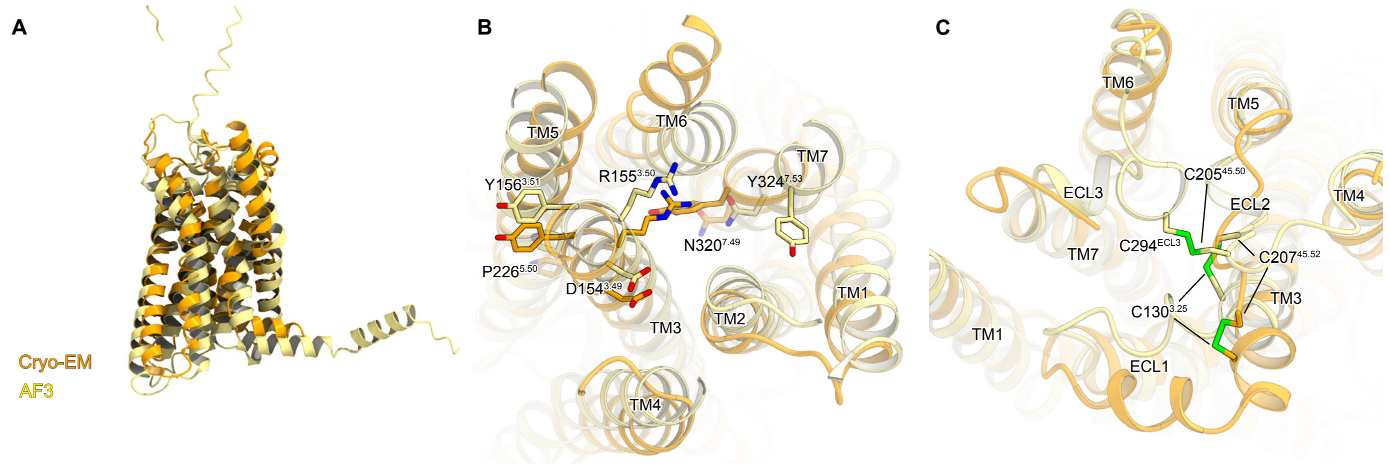


**Figure 2.—figure supplement 2. Structural comparison with AlphaFold-predicted structure.** **A-C** superimposition of the cryo-EM (orange) and AF3 (khaki) structures. (**A**) Overall view of the receptor, (**B**) focused on the intracellular side, and (**C**) focused on the extracellular side.

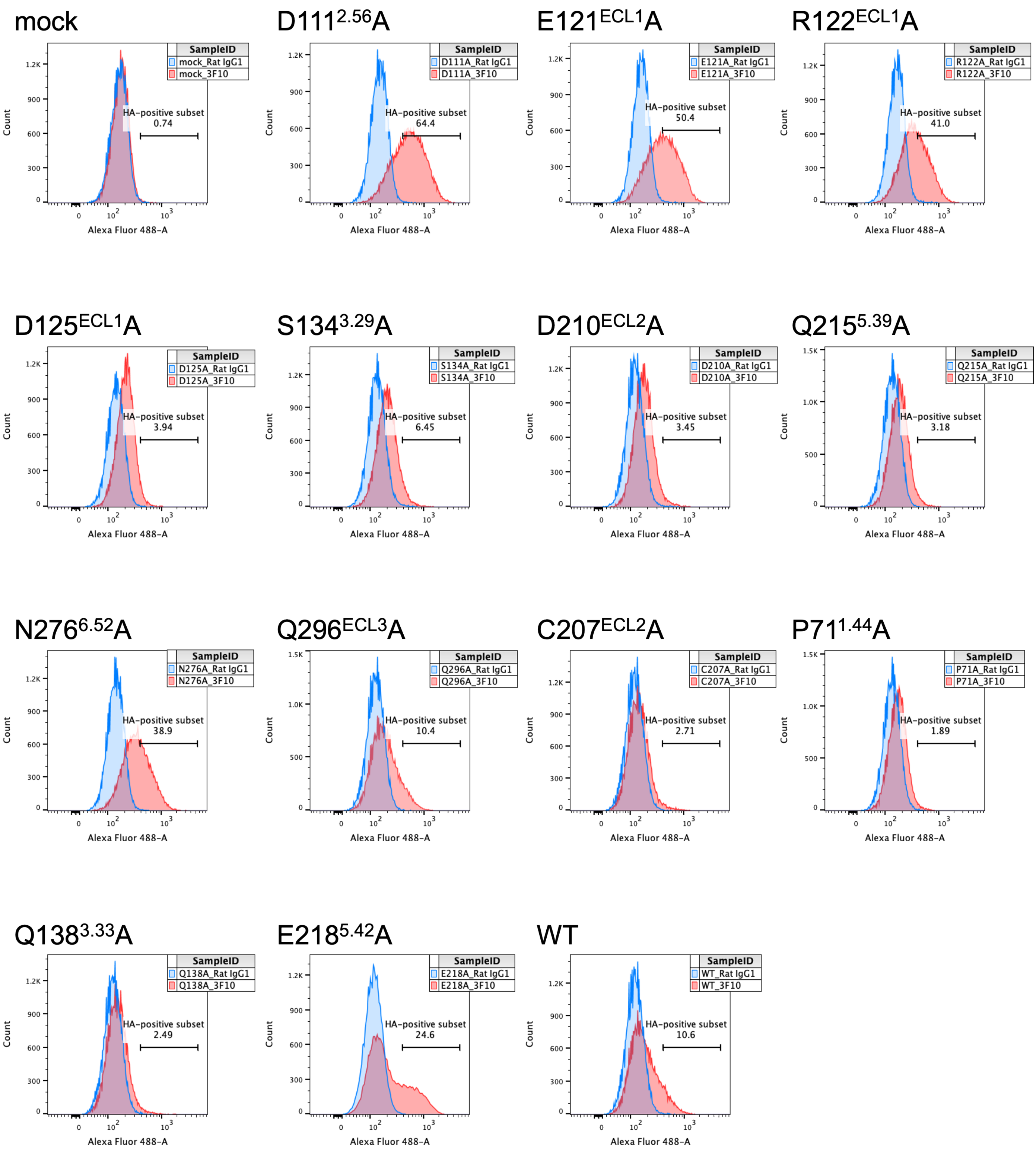


**Figure 3.—figure supplement 1 Cell surface expression of stable wild-type and GPR30 mutants.**
Flow cytometric-based assay to analyze WT and mutant GPR30 cell surface expression. Cell surface expression, defined as the HA-positive subset (red) compared with isotype control staining (blue), is indicated on each histogram. .

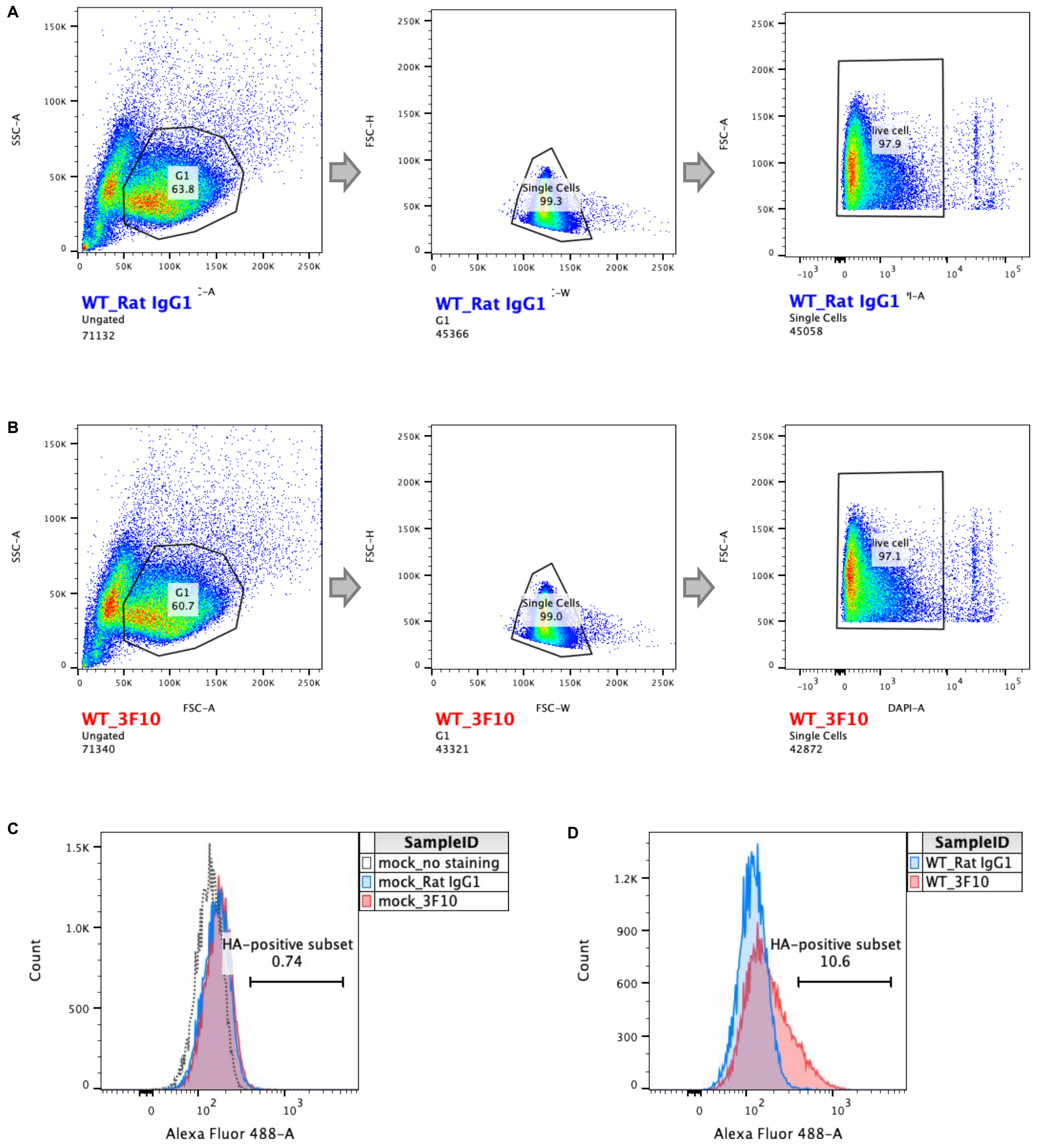


**Figure 3.—figure Supplement 2. Flow cytometric gating criteria to analyze GPR30 cell surface expression.**

**A, B** Gating used in the analysis. **C, D** Cell surface expression is defined as an HA-positive subset (red) compared with isotype control staining (blue). **(C)** Mock cells, **(D)** WT GPR30-expressing cells.

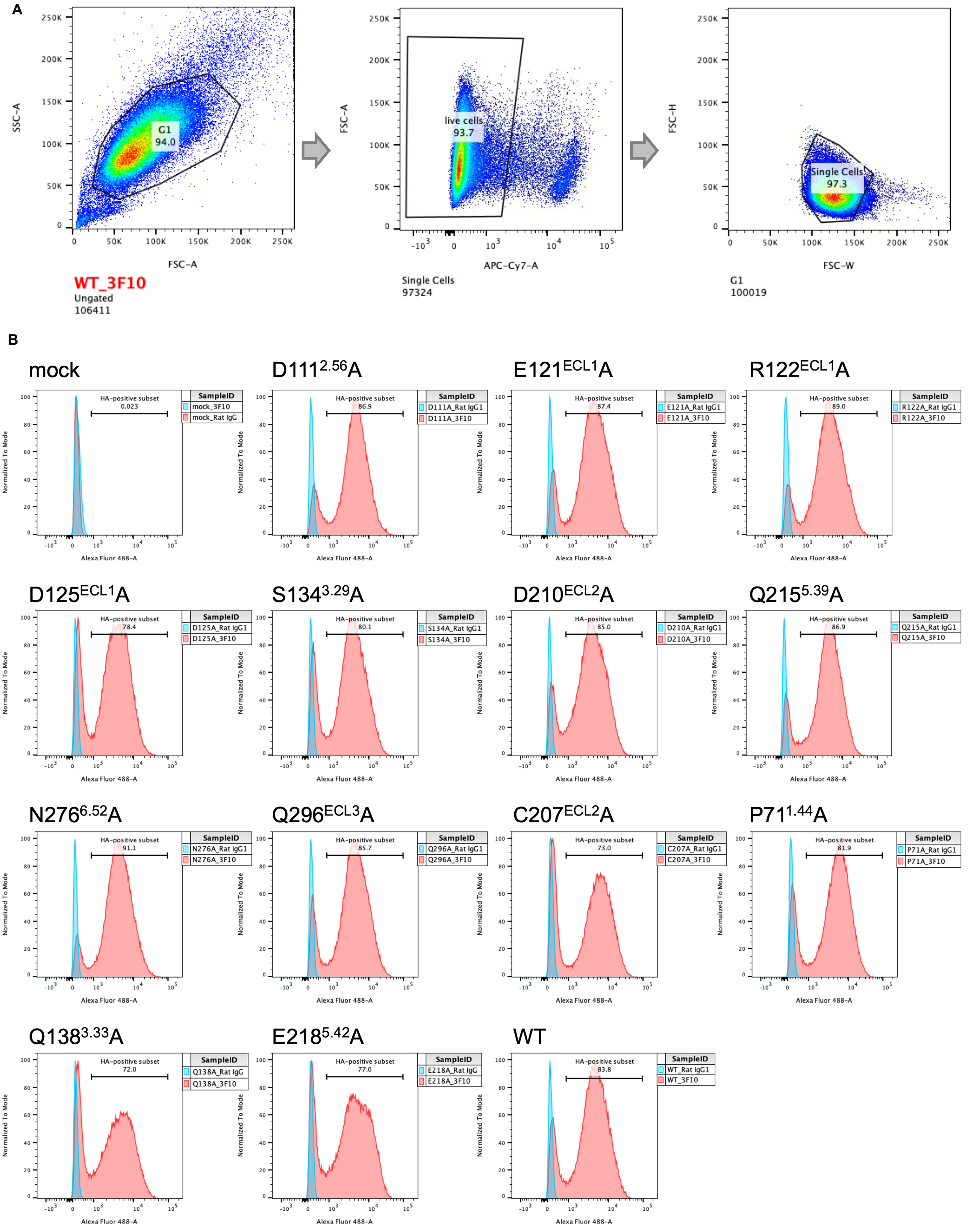


**Figure 3.—figure Supplement 3. Flow cytometric-based assay to analyze the whole-cell expression of WT and mutant GPR30.**

**A** Gating used in the analysis. **B** Whole-cell expression is defined as the HA-positive subset (red) compared with isotype control staining (blue).

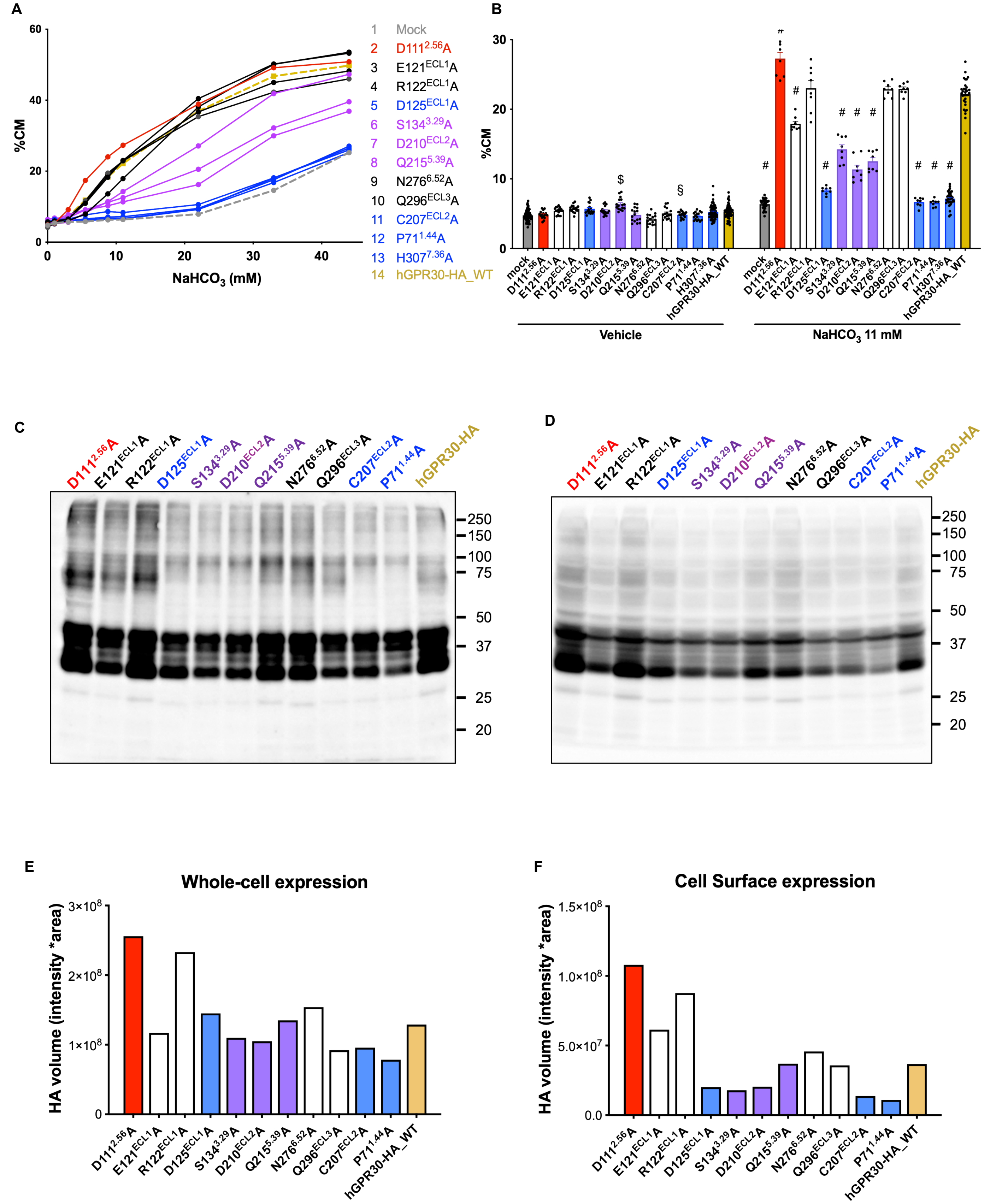


**Figure 3.—figure Supplement 4. Cell surface expression and bicarbonate-induced activation of transiently expressing GPR30 mutants.**

**A, B** TGFα shedding assay using HEK293 cells transfected with HA-tagged hGPR30. The mutants D111$^{2.56}$A; C207$^{ECL2}$A, P71$^{1.44}$A, H307$^{7.36}$A; and D125$^{ECL1}$A, S134$^{3.29}$A, D210$^{ECL2}$A, and Q215$^{5.39}$A are highlighted in red, blue, and purple, respectively. **C** Whole-cell expression of HA-tagged mutants**,** analyzed by western blotting with 20 μg of whole-cell lysate per lane. **D** Cell surface expression of HA-tagged mutants, using cell surface biotinylation and avidin immunoprecipitation, analyzed by western blotting with 1.5 μg of cell surface protein per lane. **E, F** Quantitative analysis of HA expression in **C** and **D**.

Statistical analysis: $^{\$}$ $p = 0.01$, $^{\S}$ $p < 0.001$ compared to HA-hGPR30 cells stimulated with vehicle, $^{\#}$ $p < 0.0001$ compared to HA-hGPR30 cells stimulated with 11 mM $NaHCO_3$, using two-tailed unpaired t-test with Bonferroni's correction after two-way ANOVA. Data are presented as mean values **(A)** and mean values ± SEM **(B)**.

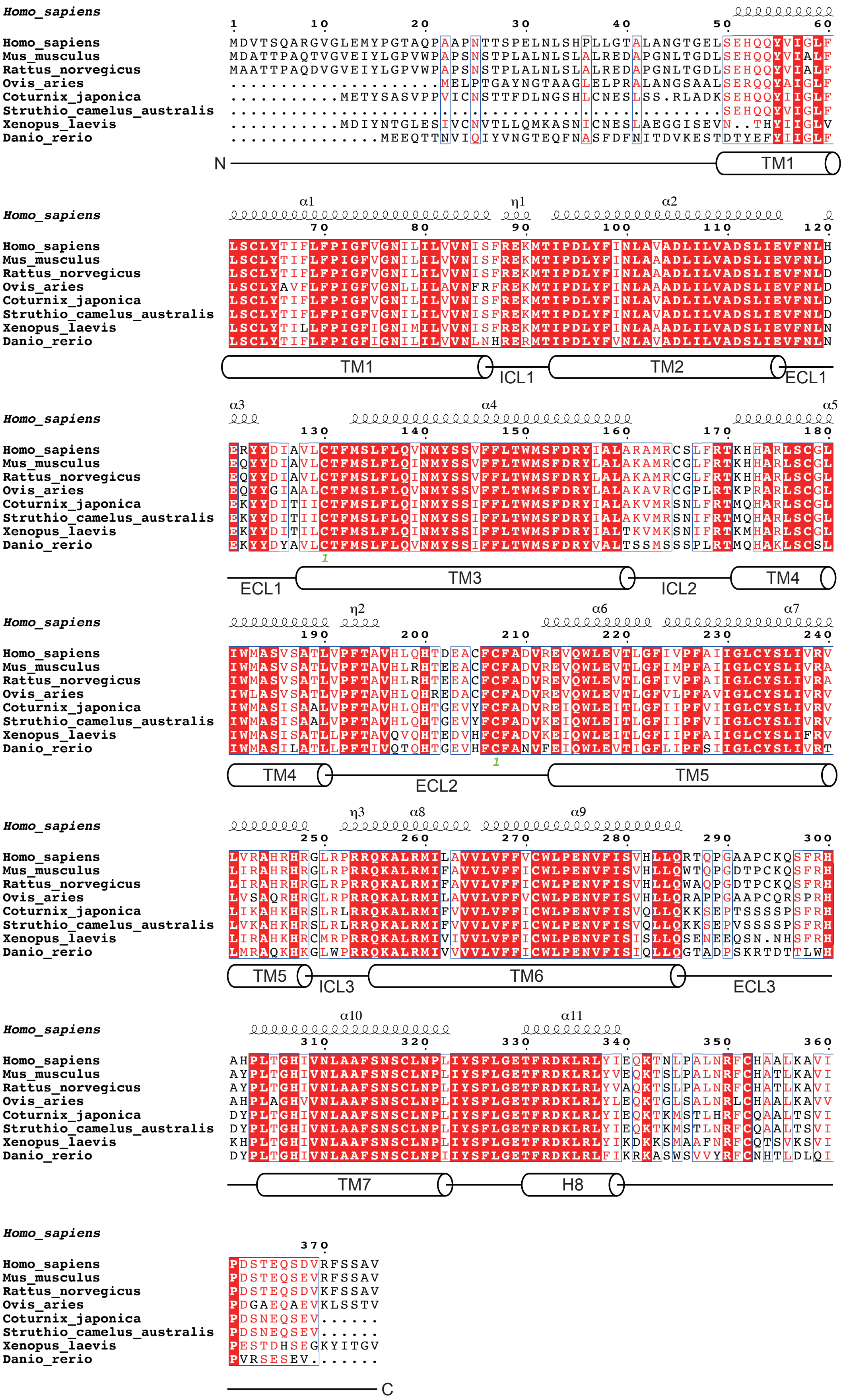


**Figure 3.—figure supplement 5.** Conservation of the GPR30 homologs. Amino acid alignment of representative GPR30 homologs.

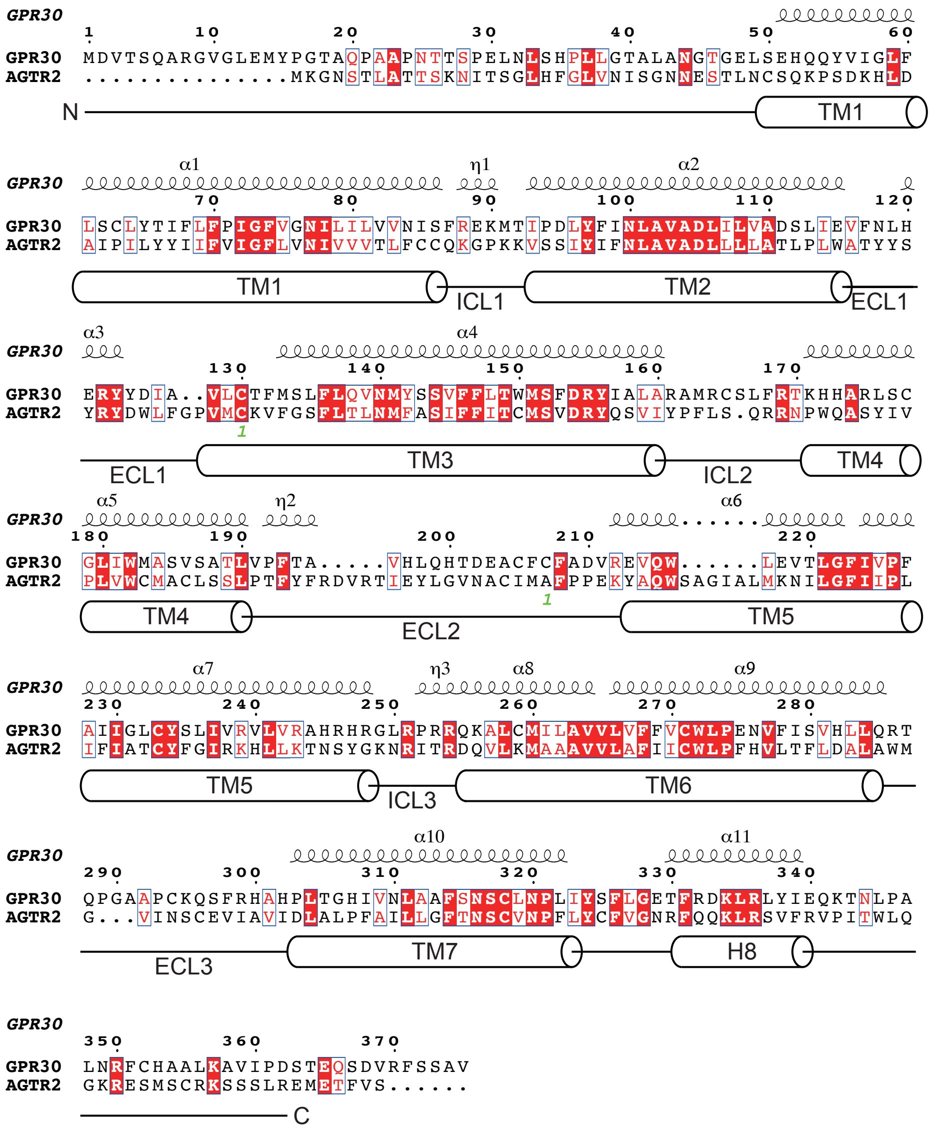

**Figure 5.—figure supplement 1**. Sequence alignment of GPR30 and AT2R.
