## supplemental table for "Cryo-EM structure of the bicarbonate receptor GPR30"

|  | | **mock** | **D111^2.56^A** | **E121^ECL1^A** | **R122^ECL1^A** | **D125^ECL1^A** | **S134^3.29^A** | **D210^ECL2^A** | **Q215^5.39^A** | **N276^6.52^A** | **Q296^ECL3^A** | **C207^ECL2^A** | **P71^1.44^A** | **Q138^3.33^A** | **E218^5.42^A** | **WT** |
| --- | --- | --- | --- | --- | --- | --- | --- | --- | --- | --- | --- | --- | --- | --- | --- | --- |
| **Best-fit values** | Top | 0.08132 | 1.244 | 0.9457 | 0.877 | 0.0868 | 0.8079 | 0.6256 | 0.8471 | 0.948 | 0.9637 | 0.06277 | 0.1079 | 0.4166 | 0.9426 | 1.155 |
|  | LogEC_50_ | 1.015 | 0.3703 | 0.6733 | 0.5393 | Unstable | 0.6966 | 0.7335 | 0.6544 | 0.4605 | 0.4903 | 1.001 | 1.216 | 1.033 | 0.5701 | 0.5838 |
|  | EC_50_ | 10.36 | 2.346 | 4.713 | 3.462 | Unstable | 4.973 | 5.414 | 4.512 | 2.887 | 3.092 | 10.02 | 16.43 | 10.79 | 3.716 | 3.835 |
| **EC50** | Null hypothesis | LogEC50 = 0.5838 | | | | | | | | | | | | | | |
|  | Alternative hypothesis | LogEC50 unconstrained | | | | | | | | | | | | | | |
|  | P value | <0.0001 | <0.0001 | 0.0391 | 0.2574 | <0.0001 | <0.0001 | <0.0001 | 0.0718 | 0.0829 | 0.1581 | <0.0001 | <0.0001 | <0.0001 | 0.8459 | 0.9996 |
| **Top (maximum response)** | Null hypothesis | Top = 1.155 | | | | | | | | | | | | | | |
|  | Alternative hypothesis | Top unconstrained | | | | | | | | | | | | | | |
|  | P value | <0.0001 | 0.0512 | 0.0249 | 0.0003 | <0.0001 | <0.0001 | <0.0001 | 0.0002 | 0.0364 | 0.0516 | <0.0001 | 0.0091 | 0.0013 | 0.106 | 0.9995 |
| **Cell surface expression (%)** | | 0.74 | 64.4 | 50.4 | 41 | 3.94 | 6.45 | 3.45 | 3.18 | 38.9 | 10.4 | 2.71 | 1.89 | 2.49 | 24.6 | 10.6 |
| **Total expression (%)** | | 0.023 | 86.9 | 87.4 | 89 | 78.4 | 80.1 | 85 | 86.9 | 91.1 | 85.7 | 73 | 81.9 | 72 | 77 | 83.8 |

**Supplementary File 1. Table of parameters calculated by nonlinear regression of the mutants’ response to bicarbonate.**

Nonlinear regression (four-parameter) was used for curve-fitting over the data shown in Figure 3F, and EC_50_ and Top (maximum response) were calculated. The values of cell surface and total expression are derived from Figure 3.—figure supplement 1 and 3, respectively.
